# Organ injury accelerates stem cell differentiation by modulating a fate-transducing lateral inhibition circuit

**DOI:** 10.1101/2024.12.29.630675

**Authors:** Erin N. Sanders, Hsuan-Te Sun, Saman Tabatabaee, Charles F. Lang, Sebastian G. van Dijk, Yu-Han Su, Andrew Labofl, Javeria Idris, Li He, Marco Marchefli, Shicong Xie, Lucy Erin O’Brien

## Abstract

To rebuild tissue form and function, injured organs accelerate the differentiation of replacement stem cell progeny. Here we demonstrate that injury-induced factors open the throttle on faster differentiation by streamlining the archetypal signaling circuit that patterns cell fates. During normal turnover of the adult Drosophila intestine, fates are patterned by a conserved lateral inhibition circuit: In stem cell pairs, mutual activation of Notch receptor by Delta ligand feeds back to create opposing states of high Notch/low Delta and low Notch/high Delta; cells terminally differentiate once their Notch activity exceeds a fate-deciding threshold. After feeding flies a gut-damaging toxin, we perform *in vivo* imaging of real-time intestinal repair and trace Notch reporter dynamics in single cells. We find that tissue damage causes the speed of Notch signal activation to accelerate dramatically; faster activation expedites terminal differentiation by propelling cells past the critical Notch threshold more quickly. Combining single-cell analyses with dynamical modeling, we show that faster activation results from aberrant elevation of Delta ligand due to loss of time-delaying circuit feedback. Injury abolishes feedback via a cytokine-JAK-STAT relay from damaged cells to stem cells, causing stem cells to deactivate the Notch co-repressor that normally turns off Delta. Thus, organ injury unmasks latent plasticity in Notch-Delta lateral inhibition to propel the differentiation of new replacement cells. By unifying temporal and spatial fate control in a single, adaptable signaling circuit, organs tune stem cell dynamics to meet environmental challenges.

## Introduction

Epithelial organs continually face environmental insults that threaten barrier integrity and demand rapid repair. To re-establish organ form and function, injury-activated stem cell divisions generate a large bolus of new cells^1–6^ while accelerated differentiation of these stem cell progeny rebuilds the epithelial barrier. Injury-accelerated differentiation has been documented in a wide diversity of epithelial barrier organs, including the mammalian and *Drosophila* intestinal tract,^2,3,7–11^ mammalian airway,^12,13^ and skin.^14,15^ Yet, while extensive studies have investigated the injury-induced signaling milieu that drives proliferation, it remains largely unknown how tissue damage accelerates differentiation.

Here we report the basis for accelerated cell differentiation following physiological organ injury in the barrier epithelium that lines the adult *Drosophila* intestine (midgut). In the fly gut, as in many mammalian organs (including airway, skin, and mammary gland), cell differentiation is controlled by Notch signaling.^12,16–25^ Midgut stem versus terminal fates are determined through an archetypal lateral inhibition circuit between equipotent stem cell pairs^26–28^: Delta ligand on one cell activates Notch on its partner; activated Notch feeds back to downregulate same-cell Delta, thereby attenuating the partner’s ability to activate the other cell’s Notch (Fig. 1a).^25,29–37^ Over time, this two-cell circuit produces a 1:1 ratio of Notch-activated and Notch-inactive cells, which in the fly midgut correspond to terminal and stem fates, respectively.

**Figure 1:**
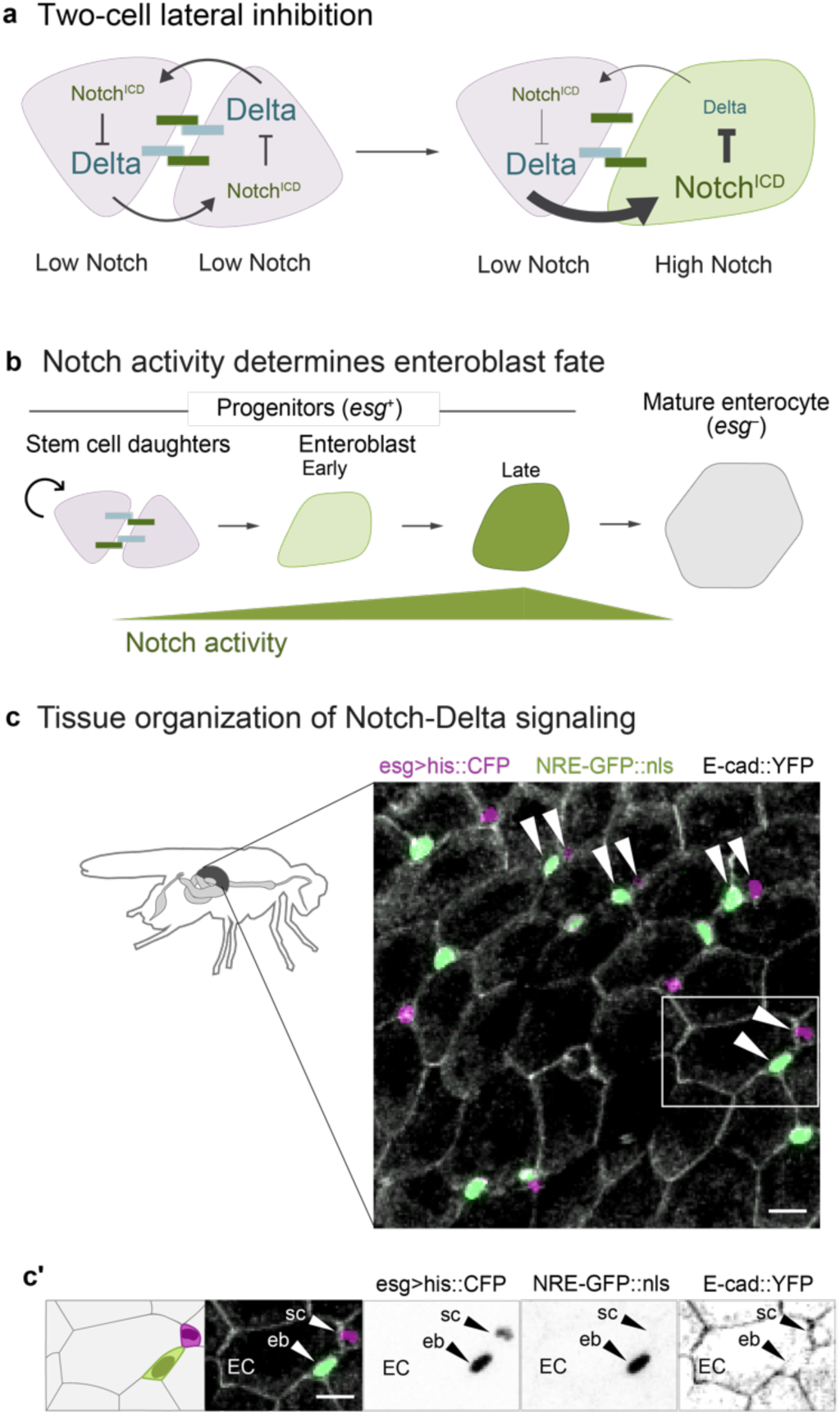
Notch-Delta signaling in the *Drosophila* adult midgut. (a) Two-cell lateral inhibition through Notch-Delta signaling. Initially, both cells express Notch receptor (dark green) and Delta ligand (blue). Stochastic differences in the two cells’ signaling levels become amplified through a feedback circuit in which Notch-Delta trans-activation and release of the Notch intracellular domain (Notch^ICD^) causes Delta to be downregulated (Fig S1a). Over time, this circuit resolves into opposing cell states of high Notch, low Delta and low Notch, high Delta. (b) Notch-Delta fate specification in the absorptive lineage. New mitotic stem cell daughters (pink) engage in mutual Notch-Delta signaling. Cell fate is determined by Notch activity: daughters that remain at sub-threshold Notch activity remain stem cells, while those that exceed the threshold differentiate into enteroblasts (early: light green; late: dark green). Enteroblasts progressively mature into terminal enterocytes (gray). The immature progenitor population (stem cells and enteroblasts) is marked by *escargot* (*esg*). (c) Tissue organization of Notch-Delta signaling. Small progenitor cells (*esg>his2b::CFP*, magenta) are interspersed among large enterocytes (outlined by *ubi-E-cad::YFP*, grayscale). Notch activity is visualized using the *NRE-GFP::nls* reporter (green; Fig S1b). Progenitors frequently form pairs of one GFP^+^ and one GFP^−^ cell (arrowheads). Both GFP^+^ and GFP^−^ cells are *esg^+^*, although GFP^+^ cells appear light green in the overlay. Scale bar, 10μm. (c’) Single-channel views of a representative cell pair (white box in c) demonstrate *esg* expression in GFP^+^ and GFP^−^ cells. Scale bars, 10μm.

We find that injury unmasks latent plasticity in this lateral inhibition circuit, turning it into a tunable mechanism to promote organ repair. Using an innovative approach for long-term, *in vivo* live imaging, we obtain an unprecedented view of native intestinal cell repair dynamics after flies have ingested a gut-damaging toxin. These movies reveal that the Notch signaling rate of differentiating cells nearly doubles, with a 200% increase in cells that accelerate signaling more than 10-fold. We further show that faster Notch signaling, and not heightened signal sensitivity, underlies faster cell differentiation.

To elucidate the basis of temporal control, we combine dynamical modeling and targeted genetic perturbations with an *in situ*, single-cell assay that quantifies the strength of lateral inhibition feedback. We demonstrate that faster signaling arises through elimination of time-delaying feedback at the core of lateral inhibition; this allows persistent expression of Delta ligand, which in turn promotes Notch receptor activation. Feedback elimination requires phospho-inactivation of Groucho (Gro), a Notch co-repressor that represses Delta transcription^28,38–40^, and is instigated by a cytokine-JAK/STAT relay from damaged cells to stem cell daughters. In this manner, the Notch-Delta fate circuit is modified by inputs from third-party cells that do not directly participate in Notch-Delta signaling. Lateral inhibition plasticity enables cells to switch from slow differentiation during healthy organ renewal to rapid differentiation for repair.

### Background: *Drosophila* midgut stem cells offer a minimal, two-cell model of Notch-Delta lateral inhibition

The *Drosophila* midgut epithelium maintains itself through continuous stem cell-driven renewal. Midgut stem cells give rise to absorptive enterocytes, which comprise >90% of differentiated cells, through a simple lineage (Fig. 1b, 1c): Stem cells both self-renew and generate enteroblasts—post-mitotic precursors that mature directly into enterocytes. These terminal enterocytes are polarized epithelial cells that establish the intestinal barrier and secrete digestive enzymes. Like in vertebrates, enterocytes in the fly gut are continually shed and replaced through stem cell division. The fly gut lacks the vertebrate intestine’s crypt-villus architecture, however. Instead, fly gut stem cells and enteroblasts are dispersed throughout the epithelium and intercalate basally among the much larger enterocytes (Fig. 1c). Together, stem cells and enteroblasts constitute the progenitor population, marked by expression of the transcription factor *escargot* (*esg*) (Fig. 1b).

The stem-versus-enteroblast fate decision distills Notch signaling to comparatively simple elements that enable detailed investigation of circuit components. In most *in vivo* models, Notch signaling is confounded by multiple Notch receptors and ligands and by more complex cellular geometries.^41–43^ By contrast, Notch signaling in the fly gut occurs between discrete progenitor cell pairs and involves exactly one Notch receptor and one ligand, Delta (Fig. 1a-c).^17,18^ Fly gut progenitors that avoid Notch activation retain stemness. Conversely, high Notch activity categorically determines absorptive fate; both Notch-null and Delta-null stem cells fail to produce enteroblasts, while ectopic Notch activation converts single stem cells into enteroblasts and, eventually, enterocytes (Fig. 1b).^16–18^

As in many systems, lateral inhibition through a canonical, Notch-Delta feedback circuit parses equipotent progenitor cells into stem and differentiated (in this case, enteroblast) fates (Fig. 1a). Signaling begins when two midgut progenitor cells make contact—either following stem cell division or through physical collision.^44^ Because activation of Notch leads to same-cell repression of Delta, small differences in Notch receptor activation between signaling cells evolve into opposing cell states of high Notch/low Delta (enteroblasts) and low Notch/high Delta (stem cells).^26–28,34^ Eventually, late-stage enteroblasts will themselves attenuate Notch as they mature into large, terminal enterocytes (Fig. 1b).^18^

Notch signal activity can be quantified using the fluorescent Notch Response Element (NRE) reporter, *NRE-GFP::nls*, that directly reflects Notch transcriptional output (Fig S1b; also known as *Su(H)Gbe* or *GBE-Su(H)*).^26,44,45^ Through live tracking of individual progenitor cells *in vivo*, we previously identified a precise *NRE-GFP::nls* intensity threshold at which midgut progenitors acquire enteroblast fate.^44^ As an individual cell progresses toward enteroblast fate, its GFP intensity increases from low to high. But when measured across the entire *esg*-expressing progenitor population, GFP intensities exhibit a consistent, bimodal distribution in both *in vivo* live imaging^44^ and fixed tissue analyses (Fig. 2a). This bimodality, with distinct low- and high-Notch peaks, reflects the binary outcome of lateral inhibition signaling. Lateral inhibition implies that cells should switch fates at the trough between low and high signaling states—a prediction we confirmed through time-resolved analysis of progenitor cells in long-term live movies.^44^ Thus, this *NRE-GFP::nls* threshold represents a critical level of Notch signaling that cells must exceed to initiate differentiation.

**Figure 2:**
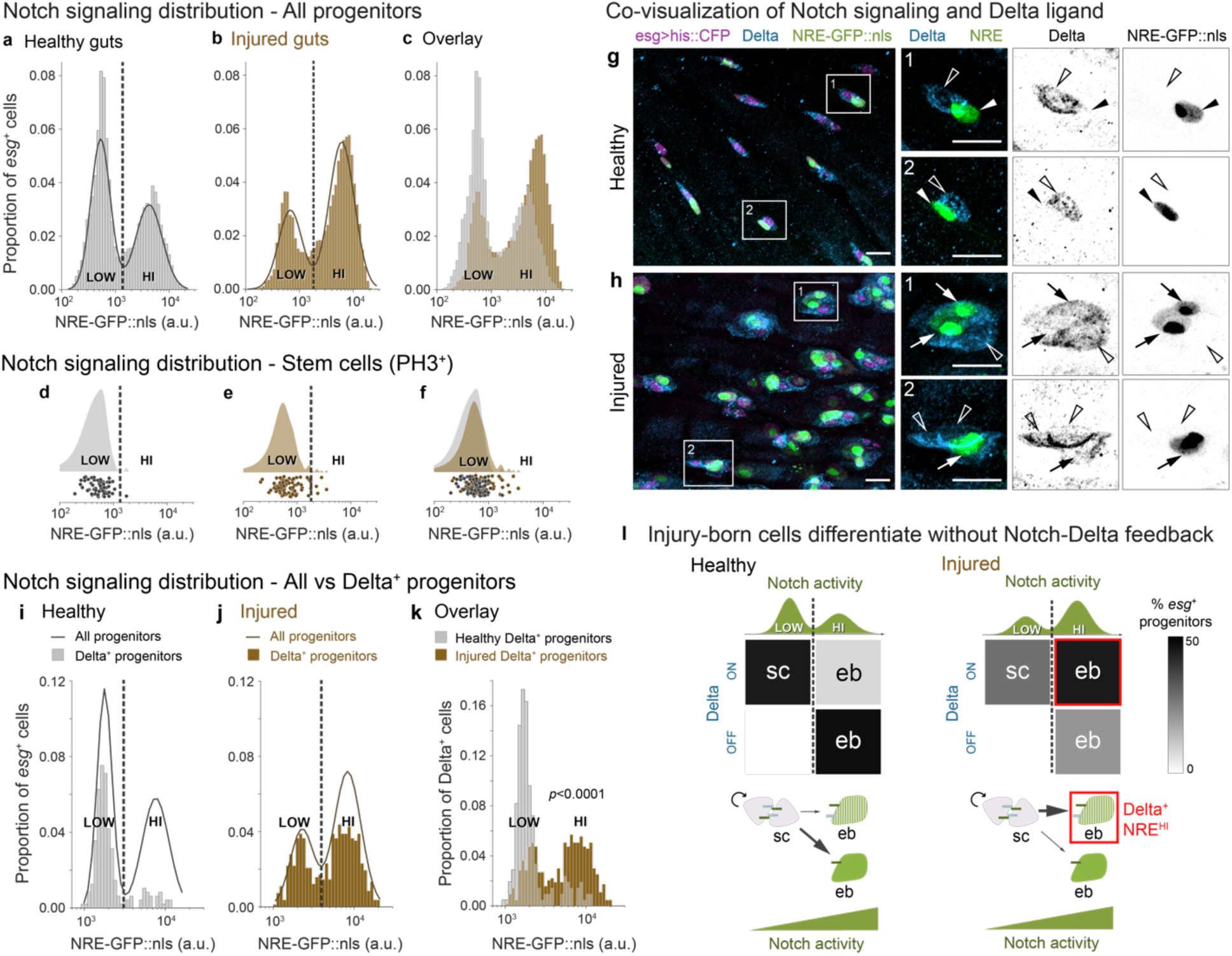
Injury disrupts Notch-Delta lateral inhibition feedback to generate Delta-expressing enteroblasts. (a-c) Notch signaling (*NRE-GFP::nls*) in progenitors (*esg>his2b::CFP*) from (a) healthy and (b) bleomycin-injured guts. Both conditions show bimodal NRE^low^ and NRE^hi^ populations (solid lines: Gaussian mixture model (GMM) fits; dashed lines: classification thresholds). (c) Overlay shows injury increases the proportion of NRE^hi^ cells while maintaining GFP intensity ranges and thresholds. Healthy: n=5681 cells, N=6 guts from a single experiment. Injury: n=8819 cells, N=6 guts from a single experiment. (d-f) *NRE-GFP::nls* in mitotic (PH3^+^) cells shown as raincloud plots (top) and single-cell measurements (bottom) from (d) healthy and (e) injured guts. Dashed lines show classification thresholds from panels a-b. In both conditions, PH3^+^ cells match the NRE^low^ peak distribution and classification (healthy: 98% NRE^low^; injured, 93% NRE^low^). (f) Overlay. Healthy: n=60 cells, N=27 guts, 7 independent replicates. Injury: n=83 cells, N=8 guts, 2 independent replicates. (As previously reported, injury sharply increases numbers of PH3^+^ cells per gut.^1,3,10,47,48,50–55^) (g-h) Co-visualization of Notch signaling (*NRE-GFP::nls*, green) and Delta immunostain (blue) in *esg>his2b::CFP* progenitors (magenta). (g) In healthy guts, Delta^+^ cells typically lack GFP and pair with Delta^-^, GFP^+^ cells. (h) In injured guts, many Delta^+^ cells show bright GFP and often form clusters with other Delta^+^, GFP^+^ as well as Delta^+^, GFP^-^ cells. Boxed regions shown at higher magnification with split channels. Arrows indicate Delta^+^, GFP^+^ cells; empty arrowheads indicate Delta^+^, GFP^-^ cells; filled arrowheads indicate Delta^-^, GFP^+^ cells. Scale bars, 10μm. (i-k) Quantification of Delta and Notch signaling relationships. Notch signaling (*NRE-GFP::nls*) specifically in Delta^+^ cells from (i) healthy and (j) injured guts, as a proportion of all *esg^+^* cells. Solid lines: GMM fits for all *esg^+^*population. *NRE-GFP::nls* raw values and classification thresholds (dashed lines) differ from panels a-c due to use of a different imaging system (see Methods). Overlay of Delta^+^ cells from (i) healthy and (j) injured guts as a proportion of Delta^+^ cells only. Injury shifts Delta^+^ cells from predominantly NRE^low^ (84%) to predominantly NRE^hi^ (62%) (*p*<0.0001). Healthy: n=478 *esg^+^* cells, n=208 Delta^+^ cells; N=2 guts from a single experiment. Injured: n=823 *esg^+^* cells, n=631 Delta^+^ cells; N=3 guts from a single experiment. *p*-value, two-sample K-S test. (l) Summary: Injury-induced disruption of Notch-Delta feedback to produce Delta-expressing enteroblasts. In healthy guts, mitotic stem cells (sc) express Delta and maintain low Notch activity, while Notch-Delta feedback drives differentiating enteroblasts (ebs) to the opposing state of high Notch activity and no Delta. In injury, differentiating enteroblasts maintain Delta despite acquiring high Notch. Gray shading indicates percent of signaling progenitors in each Notch/Delta state; green curves show GMM *NRE-GFP::nls* distributions (Fig 2a-c).

### The Notch threshold for enteroblast fate remains constant in injury

One potential mechanism to accelerate differentiation during injury would be to make the enteroblast differentiation program more sensitive to Notch signaling, such that injury-born cells acquire enteroblast fate at a lower level of Notch activity compared to healthy-born cells. Heightened sensitivity would manifest, for example, as a leftward shift in the position of the trough between NRE^low^ (stem) and NRE^hi^ (enteroblast) peaks.

To investigate this possibility, we examined the population-scale distribution of Notch signaling following injury, which we induced by feeding flies bleomycin during days 3-4 of adult life. Bleomycin is a DNA-damaging agent that targets mature enterocytes while sparing progenitor cells.^10^ At the moderate concentration (25 µg/ml) we used, barrier integrity and organismal survival are not impacted during the two-day injury period.^10^ We confirmed that bleomycin ingestion caused an increase in the per-gut number of *esg^+^* progenitor cells, as expected from damage-induced regeneration.^10,47^

We quantified *NRE-GFP::nls* intensities in individual *esg*^+^ cells from injured and healthy guts (Figs. 2a-c). The injured-gut distribution remained bimodal, with distinct low-Notch (henceforth, NRE^low^) and high-Notch (NRE^hi^) peaks. We observed that injury increased the proportion of NRE^hi^ cells (Fig. 2b), consistent with the damage-accelerated production of replacement cells by activated stem cell division.^1,3,10,47–55^ Yet, Gaussian Mixture Model (GMM) analysis revealed that, despite this shift in population distribution, injury preserves fundamental features of the NRE^low^ and NRE^hi^ states: The range of GFP intensities, spread of means, and position of the trough all remain similar (Fig. 2c). Thus, while injury shifts the proportions of low- and high-Notch states, it does not fundamentally alter the states themselves.

To determine whether the trough still marks the signaling threshold that distinguishes stem from enteroblast fate, we measured Notch activity in definitive stem cells. Mitoses are near-exclusive to stem cells in both healthy^17,18,55–58^ and bleomycin-injured guts,^55^ although their frequency increases dramatically in injury.^1,3,10,47–55^ We thus identified stem cells by immunostaining for the M-phase marker phospho-Histone H3 (PH3) and compared their *NRE-GFP::nls* intensities to the all-progenitor distributions (Fig. 2d, 2e).

We found that mitotic stem cells were overwhelmingly NRE^low^ in both healthy and injured guts (98% and 93% of PH3^+^ cells, respectively; Fig. 2d,e). Moreover, the *NRE-GFP::nls* intensity distribution of mitotic stem cells in injured guts aligned with the all-progenitor NRE^low^ population (Fig. 2b,f). These results demonstrate that stem cell identity, marked by mitotic activity, remains tightly correlated with the NRE^low^ state during injury. Since injury also preserves the NRE^low^/NRE^hi^ threshold intensity (Fig. 2c), we conclude that injury-induced enteroblast specification occurs at a similar Notch signaling threshold as in healthy conditions. Thus, injury must accelerate differentiation via mechanisms other than increased Notch sensitivity.

### Notch-Delta feedback is disrupted in injury

We next considered an alternative mechanism in which injury raises Delta ligand protein expression, which might then increase the pace of Notch-driven differentiation. To explore this scenario, we first compared the baseline relationship between Delta protein expression and Notch signaling level by Delta immunostain of healthy guts with genotype *NRE-GFP::nls, esg>his2b::CFP*. Consistent with prior reports,^17,18,26–28^ Delta^+^ cells exhibited minimal GFP and were frequently paired with Delta^−^ cells that exhibited bright GFP (Fig. 2g).

We quantified this inverse relationship by cross-correlating *NRE-GFP::nls* intensities with Delta status in single cells. The vast majority (84%) of Delta^+^ cells were NRE^low^ (Fig. 2i); a similar majority (86%) of NRE^hi^ cells were Delta^−^ (Fig. S2a, S2b). This pattern also was reflected in analyses of published single-cell transcriptomes,^59^ which revealed robust anti-correlation between Delta ligand and Notch target gene expression (Fig. S3). Altogether, these healthy-gut data exemplify lateral inhibition signaling: Below the critical Notch signal threshold (marked by the trough in the GFP distribution), cells express Delta. Although immunostaining is not a linear measure of Delta abundance, the strength of Delta expression presumably weakens as Notch signaling builds. After cells pass the critical threshold, they have largely extinguished Delta and adopt enteroblast fate.

Next, examining the injury state, we discovered a dramatic uncoupling between Delta expression and Notch activation. In injured *NRE-GFP::nls, esg>his2b::CFP* guts, immunostaining revealed numerous Delta^+^ cells with bright GFP signal (Fig. 2h), consistent with a prior report.^60^ These cells often formed clusters with other Delta^+^, GFP-expressing cells and with Delta^+^ cells that lacked GFP (Fig. 2h).

Quantifying single-cell GFP intensities, we found that 62% of Delta^+^ cells were NRE^hi^ (Fig. 2j)—a proportion four-fold greater than in healthy guts. Correspondingly, the proportion of Delta^+^, NRE^low^ cells decreased by 46% (Figs. 2i, 2j and S2b). Intriguingly, many of these Delta^+^, NRE^low^ cells were in contact with Delta^+^, NRE^hi^ cells; a recent report^61^ suggests that the Notch inhibitor Numb may enable some Delta^+^, NRE^low^ cells to escape Notch activation and thereby retain stemness. The striking emergence of Delta^+^, NRE^hi^ cells demonstrates that injury disrupts the feedback circuit that normally drives cells toward opposing signaling states (Fig. 2l).

What is the identity of this injury-induced, Delta^+^, NRE^hi^ population? Since *NRE-GFP::nls* levels distinguish cell fates even in injury—with NRE^hi^ marking enteroblasts and NRE^low^ marking stem cells (Figs. 1b and 1e)—we conclude that the Delta^+^, NRE^hi^ cells are enteroblasts that persist in expressing Delta (Fig. 2l).

### Modeling links Notch-Delta feedback to Notch signaling speed

Can persistent expression of Delta drive accelerated cell differentiation? Intriguing clues emerge from a zebrafish study in which forced Delta overexpression resulted in faster oscillations of the embryonic segmentation clock^62^ and from synthetic cell culture studies in which increasing concentrations of ectopically applied Delta resulted in faster accumulation of an engineered Notch reporter.^63,64^ To understand the relationship between Notch-Delta feedback and signaling speed, we used a mathematical model of lateral inhibition in which activation of Notch by its partner’s Delta is coupled to same-cell inhibition of Delta by activated Notch (Fig. 3a; see Methods).^34^ The model is governed by two dimensionless parameters: K_N_, which is the threshold for Notch activation by Delta, and K_D_, which is the threshold for Delta inhibition by Notch (Fig. 3a). Both cells initially have high Delta and low Notch, with symmetry broken by a slight elevation of Notch in one cell. The time evolution of Notch activity and Delta level is defined by Eqs. 1 and 2 (Fig. 3a) using experimentally derived parameter ranges from healthy guts (see Methods).^27^

**Figure 3:**
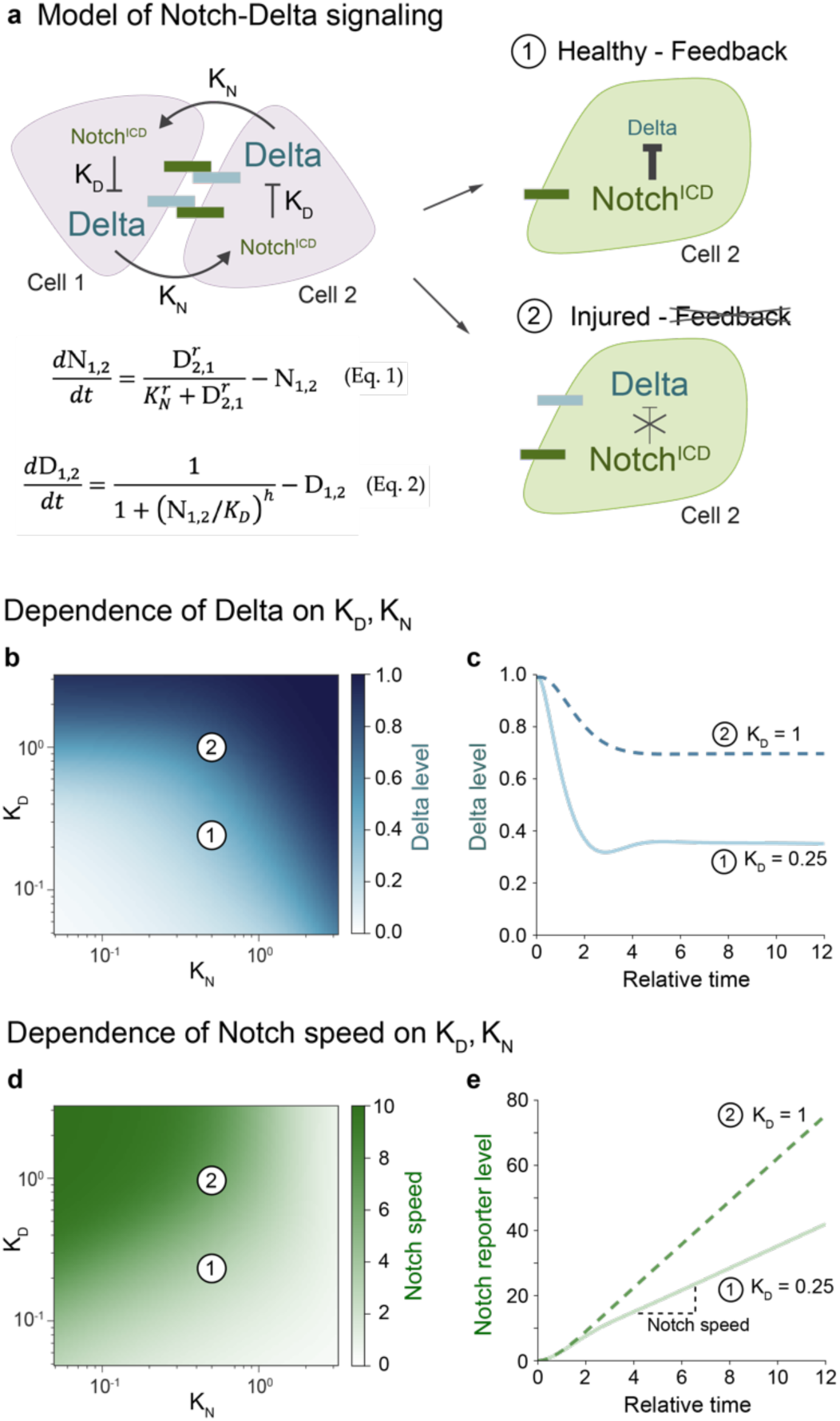
Disrupted Notch-Delta feedback can accelerate Notch signaling. (a) Model schematic for Notch-Delta lateral inhibition.^27,34^ Two key parameters govern the system: K_N_ (the threshold for Notch activation by Delta) and K_D_ (the threshold for Delta inhibition by Notch). Cell 2 is initialized with slightly higher Notch activity. Outcomes 1 (high-Notch/low-Delta) and 2 (high-Notch/high-Delta) represent the dominant enteroblast states in healthy and injured guts, respectively. Equations 1-2 describe the time evolution of Notch activity and Delta levels. Hill coefficients r=h=2. (b-e) Model parameter space and dynamics. Parameter values for Point 1 (K_N_=0.5, K_D_=0.25); Point 2 (K_N_=0.5, K_D_=1). (b) Steady-state Delta level (t=10) as a function of K_N_ and K_D_. While injury decreases K_N_ and increases K_D_ (see Results), only increased K_D_ reproduces the high-Notch/high-Delta injury state. (c) Simulated time evolution of Delta levels for Points 1 and 2. See Fig. S4a for additional K_D_ values. (d) Notch signaling speed as a function of K_N_ and K_D_. Signaling speed is defined as the mean rate of Notch reporter accumulation from t=4 to t=10. Increased K_D_ accelerates signaling speed. (e) Simulated time evolution of Notch reporter levels for Points 1 and 2. See Fig. S4b for additional K_D_ values.

Since K_N_ is inversely proportional to cell-cell contact area,^27^ and contact area increases in injury due to both cell enlargement and clustering (compare Fig. 2g,h),^60^ we first investigated whether decreased K_N_ could explain the injury-induced high-Notch/high-Delta state. Reducing K_N_ in our simulations failed to reproduce the injury phenotype, however; instead of maintaining high Delta, cells with high Notch showed reduced Delta (Fig. 3b). This outcome persisted in a three-cell model simulating injury-induced clusters (see Modeling Supplement). Thus, changes in K_N_ alone cannot account for the injury phenotype.

We then examined K_D_, which inversely relates to Notch’s ability to suppress Delta. The persistence of Delta expression in high-Notch cells during injury suggested elevated K_D_. Indeed, increasing K_D_ in both two- and three-cell simulations generated high-Notch cells with elevated Delta, reproducing the injury state (Fig. 3b, 3c; Modeling Supplement).

Having identified increased K_D_ as the key parameter change, we next investigated its effect on Notch signal dynamics. When K_D_ is elevated, signaling cells maintain higher Delta levels and thus provide more ligand to activate Notch. We hypothesized that increased Delta would accelerate Notch target gene accumulation and thus cell differentiation. To test this prediction, we added a Notch-driven reporter to our model (see Methods) and calculated Notch signaling speed as the rate of reporter accumulation. Consistent with our hypothesis, increased K_D_ led to faster Notch signaling during the initial, linear phase of signaling across a broad range of values for K_N_ (Fig. 4d, e). Overall, these simulations align with prior overexpression studies^62–64^ and predict that injury-induced disruption of Notch-Delta feedback accelerates Notch signaling speed.

**Figure 4:**
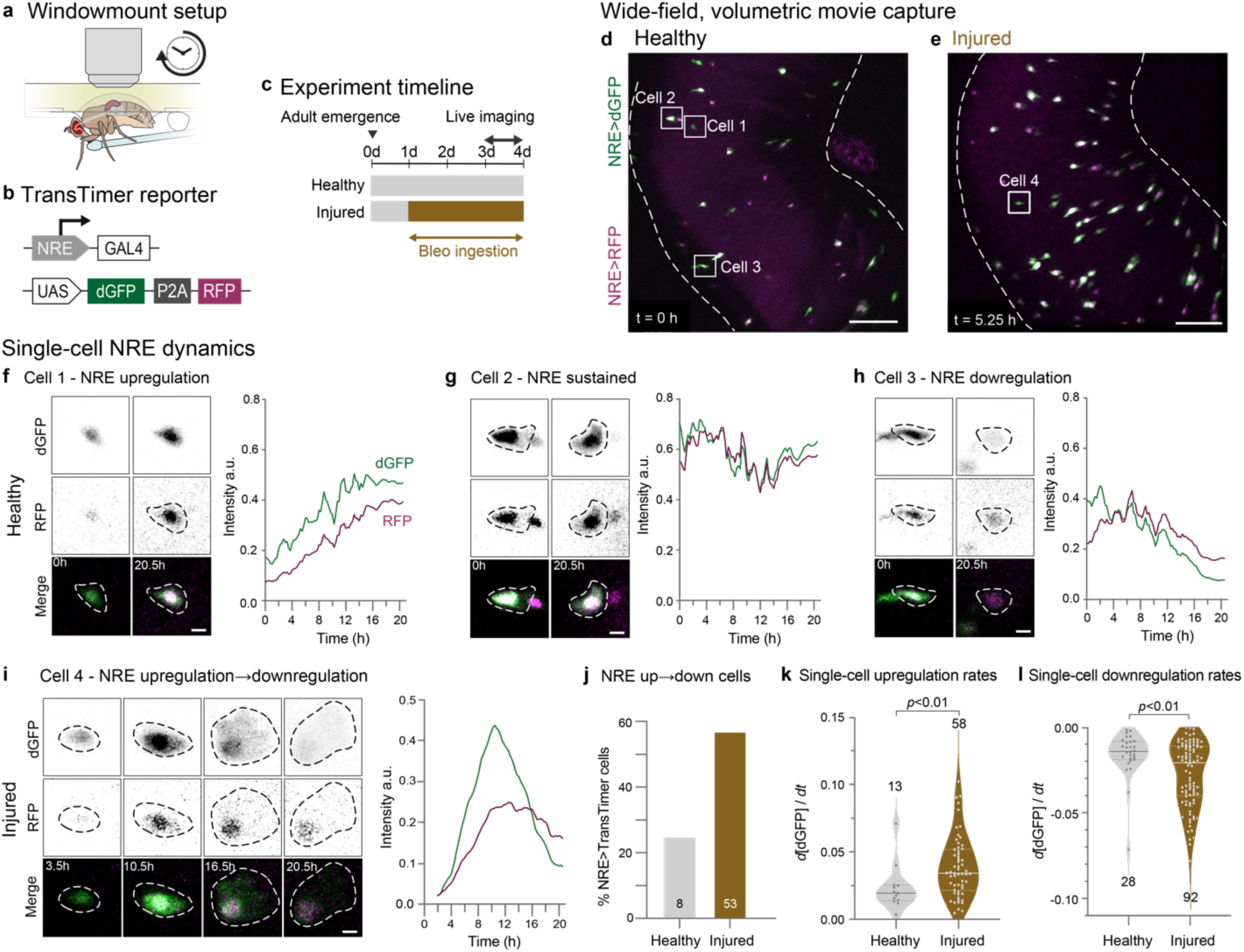
Real-time *in vivo* imaging reveals acceleration of Notch signaling dynamics in differentiating cells during injury. (a) Windowmount setup for long-term imaging of actively feeding flies.^44^ A window cut in the dorsal cuticle enables the intact gut to be imaged in a near-native context for >20 hours. Flies receive nutrients and bleomycin (for injury condition) through a microcapillary feeder tube. (b) Design of the NRE>TransTimer dual-color kinetic reporter. NRE-GAL4 (Notch Response Element: *c.f*. Fig S1b) drives expression of UAS-TransTimer^65^: a bicistronic cassette that encodes fast-folding, destabilized dGFP and slow-folding, stable RFP, separated by P2A. (c) Experimental timeline. Prior to imaging, flies are fed standard fly food either with or without 25 µg/mL bleomycin in yeast paste. During imaging, flies are fed 10% sucrose with or without 25 µg/mL bleomycin. (d-e) Wide-field, volumetric live imaging of NRE>TransTimer guts in (d) healthy and (e) injured conditions. Images are stack projections of single timepoints from 20.5-hour Windowmount movies. Dotted lines indicate gut boundaries. Individual NRE>TransTimer cells exhibit specific dGFP/RFP ratios that indicate distinct stages of differentiation. Numbered boxes mark Cells 1-4 analyzed in Panels f-i. Scale bars: 50 µm. See Movies 1 and 2. (f-i) Real-time dynamics of NRE>TransTimer in single differentiating cells. Images are zoomedin stack projections showing single- and two-channel views of Cells 1-4 at the timepoints indicated. Corresponding traces show movie-normalized dGFP and RFP intensities over time (see Methods). In healthy-gut Movie 1, Cells 1-3 each exhibit a distinct phase of differentiation: NRE upregulation, sustained signaling, or downregulation. In injured-gut Movie 2, Cell 4 progresses through all phases within the same 20.5-hour imaging period. Scale bars: 5 µm. See Movies 3-6. (j) Injury accelerates the differentiation-associated Notch signaling program. During 20.5-hour movies, cells exhibiting both up- and downregulation of NRE>TransTimer are 2.2.-fold more abundant in injured guts than healthy guts (57% versus 25%, respectively). Data from N=2 healthy-gut movies (n=8/32 total cells) and N=3 injured-gut movies (n=53/93 total cells). (k-l) Injury increases real-time rates of single-cell Notch signaling. Each dot shows rate of dGFP intensity change during phases of (k) NRE>TransTimer upregulation and (l) downregulation. Medians: upregulation (injured: 0.034, healthy: 0.019), downregulation (injured: 0.021, healthy: 0.014). Data from N=2 healthy-gut movies (n=13 upregulating, 28 downregulating cells) and N=3 injured-gut movies (n=58 upregulating, 92 downregulating cells). Box plots show medians and quartiles. *p*-values from Mann-Whitney test.

### Notch signaling accelerates during injury

To test our model prediction, we performed *in vivo* live imaging of single-cell Notch dynamics in healthy and injured guts. If injury accelerates Notch signaling speed, we would expect two observable outcomes. First, live reporters of Notch activity should accumulate more rapidly in injured guts compared to healthy controls. Second, since Notch signaling naturally turns off during late stages of terminal maturation (Fig. 1b), individual differentiating cells should spend less total time in the NRE^hi^ state as they are propelled more quickly through the differentiation process.

Using the Windowmount protocol,^44^ we created a viewing window in the animal’s dorsal cuticle (Fig. 4a), enabling midgut imaging in awake, moving flies. Unlike *ex vivo* approaches, Windowmount preserves complete GI anatomy and physiology: The GI tract and its associated tissues––neurons, trachea, immune cells, and fat body––remain intact and functional. Flies continue to ingest food, undergo peristalsis, and defecate throughout imaging sessions that last over 20 hours.^44^ Consequently, Windowmount movies vividly capture midgut cell dynamics in a near-native physiological context.

To monitor Notch signaling in individual cells with high temporal resolution, we used a dual-color kinetic reporter (UAS-TransTimer)^65^ driven by the Notch Response Element (NRE-GAL4)^66^ (Fig. 4b). The TransTimer’s fast-folding, destabilized GFP (dGFP – maturation ∼0.1 h; half-life ∼2h)^65^ sensitively reports changes in NRE-GAL4 activity. Its slow-folding, long-lived RFP (maturation ∼1.5 h; half-life ∼20 h)^65^ enables identification of cells that have recently deactivated Notch (Fig. 1b) as RFP-only.

We acquired two-channel Windowmount movies of NRE-driven TransTimer (hereafter NRE>TransTimer) in healthy and injured midguts of 3-day-old adults (Fig. 4d, 4e; Movies 1-2). Three key elements enabled us to generate granular data of high quality: (1) organ-scale, volumetric imaging (∼250x250x150 µm) for unbiased, 4D capture of multiple NRE>TransTimer cells per gut; (2) micron-resolution for precise single-cell segmentation; and (3) high temporal sampling (7.5-minute intervals) over 20-hour sessions—a timeframe compatible with Notch signal dynamics.^44^ We tracked individual NRE>TransTimer cells from their first appearance until either signal loss or the movie’s end. At each timepoint, we quantified single-cell GFP and RFP intensities to generate traces of real-time NRE dynamics (see Methods).

Previous analyses imply that Notch-driven differentiation proceeds through distinct phases of signal upregulation, stability, and downregulation (Fig. 1b).^16,17,44^ Using NRE>TransTimer, we directly observed and analyzed these phases in real-time (Fig. 4f-i; Movies 3-6). While cell traces from both healthy and injured guts exhibited all three phases, their dynamics differed dramatically: In healthy-gut movies, most cells (75%) exhibited just one signaling phase over 20.5 hours of imaging, with only 25% of cells exhibiting both up- and downregulation, consistent with reported rates of organ-scale turnover.^11,47,67^ In injured-gut movies, by contrast, most cells (55%) exhibited both up- and downregulation within this same timeframe—a 2.2-fold increase over healthy guts. These data demonstrate that many differentiating cells complete the full Notch signaling program within one day during injury-induced regeneration, compared to multiple days during normal turnover. Thus, injury accelerates the temporal dynamics of the Notch signaling program itself in addition to the overall rate of cell differentiation.

To quantify the magnitude of these dynamics, we measured rates of NRE up- and downregulation using the time-resolved measurements of fast-folding, fast-degrading dGFP (see Methods). In up-down traces, clear temporal separation was evident between fast-folding, destabilized dGFP and slow-folding, stable RFP, matching the expected reporter dynamics.^65^ In traces that were up-only or down-only, minimal or no temporal offset was observed, indicating that signal dynamics occurred on timescales similar to or slower than RFP’s decay kinetics. The presence of these distinct dGFP/RFP patterns—even among neighboring cells in the same field of view—underscores a crucial principle: accurate interpretation of TransTimer ratios in fixed samples requires careful consideration of how signal duration compares to reporter kinetics.

Analyses of TransTimer dGFP revealed that injury increased the median rate of NRE upregulation by 75% (median d[dGFP]/dt = 0.034 versus 0.020 a.u. in healthy-gut movies). Moreover, the proportion of cells that exhibited rapid activation (defined as d[dGFP]/dt>0.025) rose from 23% in healthy guts to 69% in injured guts (Fig. 4k). Similarly, NRE downregulation was 69% faster in injury (mean d[dGFP]/dt = -0.027 versus - 0.016 a.u. in healthy guts), with cells showing rapid deactivation increasing from 7% to 47% (Fig. 4l).

The direct demonstration of accelerated Notch signal dynamics confirms our model’s prediction that disrupting Notch-Delta feedback enhances Notch signaling speed (Fig. 3d and 3e). Together, these theoretical and experimental findings implicate injury-induced modulation of lateral inhibition circuitry as accelerating the transduction of the fate-determining Notch signal during regeneration.

### Groucho inactivation underlies injury-induced feedback disruption

How does injury result in elevated K_D_? An appealing mediator is the Notch-modulated co-repressor Groucho (Gro), a global transcriptional corepressor that acts to regulate Notch signal transduction (Fig. S1a).^38,68^ Following Notch activation, Gro interacts with Notch-induced E(spl)-C proteins to suppress Delta expression.^28,40^ Hence, Gro is ideally situated to modulate K_D_, with consequent effects on Notch-Delta feedback. Supporting this notion, depletion of Gro in *Drosophila* midgut progenitors generates cells that simultaneously exhibit Delta expression and Notch activity^28^ (Fig. 5a)—a phenotype reminiscent of the Delta^+^, NRE^hi^ cells we observe in injury.

**Figure 5:**
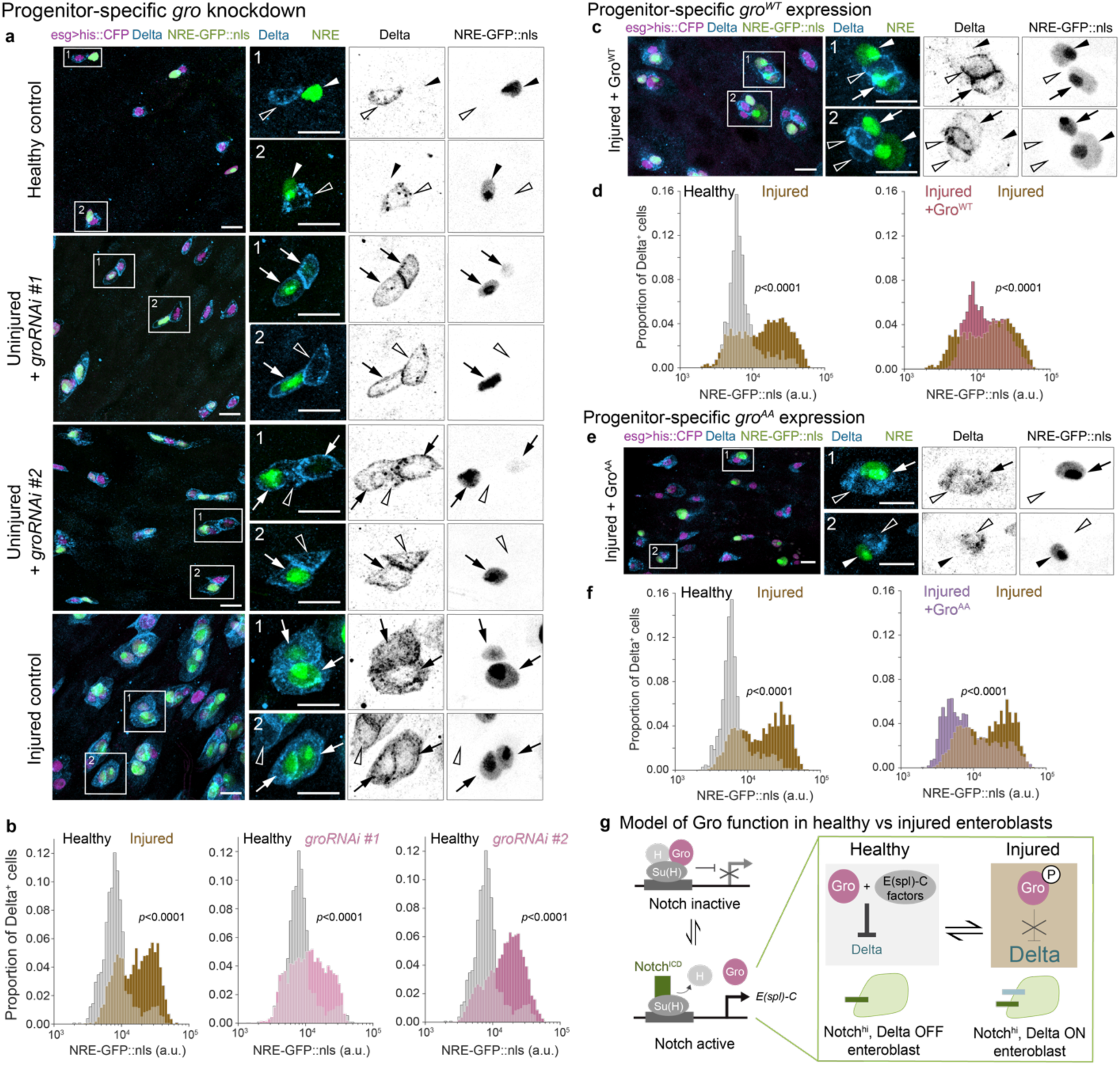
Gro inactivation attenuates Notch-Delta feedback in injury. (a) Co-visualization of Notch signaling (*NRE-GFP::nls*, green) and Delta immunostain (blue) in progenitor cells (*esg^ts^>his2b::CFP*, magenta) from healthy control guts, uninjured guts with *groRNAi* (*esg^ts^>his2b::CFP, groRNAi*), and injured control guts. Two RNAi constructs were used: *groRNAi* #1 – VDRC #KK110546; *groRNAi* #2 – BDSC #91407. In healthy control guts, cells express either GFP or Delta but rarely both. Knockdown of *gro* in uninjured guts causes cells to express both markers, mimicking the injury state. Boxed regions shown at higher magnification with split channels. Arrows indicate Delta^+^, GFP^+^ cells; empty arrowheads indicate Delta^+^, GFP^-^ cells; filled arrowheads indicate Delta^-^, GFP^+^ cells. Scale bars, 10μm. (b) Notch signaling distributions (*NRE-GFP::nls*) of Delta^+^ progenitors from the same genotypes and conditions in panel a. In otherwise uninjured guts, *esg^ts^>groRNAi* increases the proportion of NRE^hi^ cells (*groRNAi* #1 - 32%, #2 - 51%) compared to healthy controls (11%), reaching levels similar to injured controls (46%). Healthy control: n=1328 cells; N=7 guts. Injured control: n=2251 cells; N=6 guts. Uninjured *groRNAi* #1: n=4766 cells; N=14 guts. Uninjured *groRNAi* #2: n=6945 cells; N=14 guts. Samples are from 3 independent experiments. *p*-values from two-sample K-S test. See also Figures S2c, S5b. (c) Co-visualization of *NRE-GFP::nls* (green) and Delta immunostain (blue) in *gro^WT^*-expressing progenitor cells from injured guts (*esg^ts^>his2b::CFP, UAS-gro^WT^*, magenta). Although some progenitors express Delta and not GFP, or GFP and not Delta, many still co-express these two markers. Arrows indicate Delta^+^, GFP^+^ cells; empty arrowheads indicate Delta^+^, GFP^-^ cells; filled arrowheads indicate Delta^-^, GFP^+^ cells. Scale bars: 10 µm. (d) *NRE-GFP::nls* distributions of Delta^+^ progenitors from healthy control guts, injured control guts, and injured *esg^ts^>gro^WT^*guts. In injured *esg^ts^>gro^WT^* guts, the proportion of NRE^hi^ cells remains elevated (58% - injured *gro^WT^*, 62% - injured control) compared to healthy controls (15%). Healthy control: n=821 cells; N=7 guts. Injured control: n=2814 cells; N=5 guts. Injured *gro^WT^*: n=738 cells; N = 11 guts. Samples are from 3 independent experiments. *p*-values from two-sample K-S test. See also Figures S2d, S5c. (e) Co-visualization of *NRE-GFP::nls* (green) and Delta immunostain (blue) in injured-gut progenitor cells expressing constitutively active Gro^AA^ (*esg^ts^>his2b::CFP, UAS-gro^AA^*, magenta). Many progenitors exhibit either only GFP or only Delta, akin to healthy controls. Few cells still co-express Delta and GFP. Arrows indicate Delta^+^, GFP^+^ cells; empty arrowheads indicate Delta^+^, GFP^-^ cells; filled arrowheads indicate Delta^-^, GFP^+^ cells. Scale bars: 10 µm. (f) *NRE-GFP::nls* distributions of Delta^+^ progenitors from healthy control guts, injured control guts, and injured *esg^ts^>gro^AA^*guts. In injured guts, Gro^AA^ reduces the proportion of NRE^hi^ cells (27% - injured *gro^AA^*; 55% - injured control), approaching that of healthy controls (19%). Healthy control: n=1083 cells; N=5 guts. Injured control: n=2581 cells; N=5 guts. Injured *gro^AA^*: n=2119 cells; N=11 guts. Samples are from 3 independent experiments. *p*-values from two-sample K-S test. See also Figures S2e, S5d. (g) Schematic of Gro function in healthy versus injury-disrupted lateral inhibition. In the absence of Notch activation (Notch OFF), Gro associates with Su(H) and H at Su(H) binding elements to repress Notch target genes including the *E(spl)-C*.^38,68^ Upon Notch activation (Notch ON), Notch^ICD^ displaces Gro and H, binding Su(H) to drive *E(spl)-C* expression.^40^ In healthy guts, released Gro then partners with Notch-induced *E(spl)-C* factors to repress Delta. In injured guts, by contrast, phospho-inactivation of Gro precludes its association with *E(spl)-C* factors, allowing sustained Delta expression in Notch-activated cells.

To quantify the effect of *gro* depletion on Notch-Delta feedback, we performed *tub-GAL80^ts^*-inducible knockdown of *esgGAL4*-driven *groRNAi* (*esg^ts^>groRNAi*), using two independent RNAi lines. Focusing on Delta^+^ progenitors, we measured single-cell *NRE-GFP::nls* intensities and compared the resulting GFP distributions to those from healthy and injured guts of control genotype (Fig. 5b). As expected, injured control guts showed a marked increase in Delta^+^ cells that were NRE^hi^ (Fig. 5b: 11%-healthy, 46%-injured). Notably, depleting *gro* in otherwise uninjured guts resulted in Delta^+^ cells exhibiting NRE^hi^ proportions similar to the injured controls (Fig 5b: 32%-*groRNAi* #1, 51%-*groRNAi* #2). Similarly, the proportion of NRE^hi^ cells that were Delta^+^ increased dramatically in *esg^ts^>groRNAi* guts (Figs. S2c, S5b: 83%-*groRNAi* #1, 88%-*groRNAi* #2), surpassing both healthy and injured controls (Fig. S2c: 19%-healthy, 59%-injured). These results corroborate that Gro is essential to suppress Delta during Notch-Delta feedback and suggest that injury may disrupt this feedback by inhibiting Gro.

To test whether Gro inactivation is responsible for Notch-Delta feedback disruption during injury, we asked whether blocking this inactivation can re-establish Delta suppression in Notch-activated cells during injury. We performed conditional, progenitor-specific expression using two distinct *gro* alleles: wild-type *gro^WT^* (*esg^ts^>gro^WT^*) and constitutively active *gro^AA^* (*esg^ts^>gro^AA^*), which contains non-phosphorylatable mutations (T308A/S510A) that block its inactivation.^69^ We then analyzed Notch-Delta feedback using *NRE-GFP::nls* intensities and Delta immunostaining.

Wild-type and constitutively active Gro showed distinct capacities to restore normal Notch-Delta feedback after injury. In injured *esg^ts^>gro^WT^* guts, rescue was limited: Although NRE^hi^ cells that were Delta^+^ decreased sharply (Figs. S2d, S5c; 39%-injured gro^WT^, 75% - injured controls), most Delta^+^ cells remained NRE^hi^ (Fig. 5d; 58% - injured *gro^WT^*, 62% - injured control), and many *gro^WT^* progenitors continued to co-express Delta and GFP (Fig. 5c). By contrast, constitutively active *gro^AA^* achieved a robust rescue: In injured *gro^AA^* progenitors, Delta and GFP were often mutually exclusive (Fig. 5e), akin to healthy controls. Indeed, *esg^ts^>gro^AA^* guts showed substantial reductions in both NRE^hi^ cells that were Delta^+^ (Figs. S2e and S5d; 38% - *gro^AA^*, 62% - injured control) and in Delta^+^ cells that were NRE^hi^ (Fig. 5f; 27% - *gro^AA^*; 55% - injured control). The graded rescue of Notch-Delta feedback in injured guts—partial with *gro^WT^*, robust with *gro^AA^*—demonstrates that ectopic Gro activity can override Notch-Delta feedback disruption in injured guts.

Together, these loss- and gain-of function phenotypes support a model in which injury inactivates Gro. Deprived of the Gro co-repressor activity, injury-born enteroblasts fail to suppress Delta, resulting in elevated K_D_ and consequently faster Notch signaling speed.

### Injury-responsive activation of Domeless-JAK-STAT relays a damage signal to progenitors

Finally, we investigated the tissue-level injury signals that prompt cells to disrupt Notch-Delta feedback. A prime candidate is the JAK-STAT pathway, whose role in promoting regeneration is widely conserved across numerous metazoan tissues.^70–74^ In the *Drosophila* midgut, damaged enterocytes secrete cytokines that activate JAK-STAT signaling in progenitor cells by binding to the Domeless receptor.^2,47,48,52,53,60,72,75–77^ Although the Domeless-JAK-STAT pathway is primarily known for stimulating stem cell proliferation during injury, its activation in both stem cells and enteroblasts^2,47,51^ suggests it may also serve as the injury signal that disrupts Notch-Delta feedback.

To examine this possibility, we blocked JAK-STAT by conditionally expressing a dominant negative allele of *domeless* (*dome^DN^*)^78^ in progenitors (*esg^ts^>dome^DN^*). Strikingly, *dome^DN^* effectively restored Notch-Delta feedback to injured guts at near-healthy levels. *dome^DN^* progenitors typically exhibited either Delta or *NRE-GFP::nls*, but rarely both (Fig. 6a). The proportion of Delta^+^ cells that were NRE^hi^ dropped from 54% in injured controls to 14% in injured *esg^ts^>dome^DN^* guts, nearly matching healthy controls (11%) (Fig. 6b). Similarly, the proportion of NRE^hi^ cells that were Delta^+^ decreased from 69% in injured controls to 10% in injured *esg>dome^DN^* guts, surpassing healthy controls (16%) (Fig. S2f, S5e). These results establish Domeless-JAK-STAT as essential for injury-induced disruption of Notch-Delta feedback.

**Figure 6.**
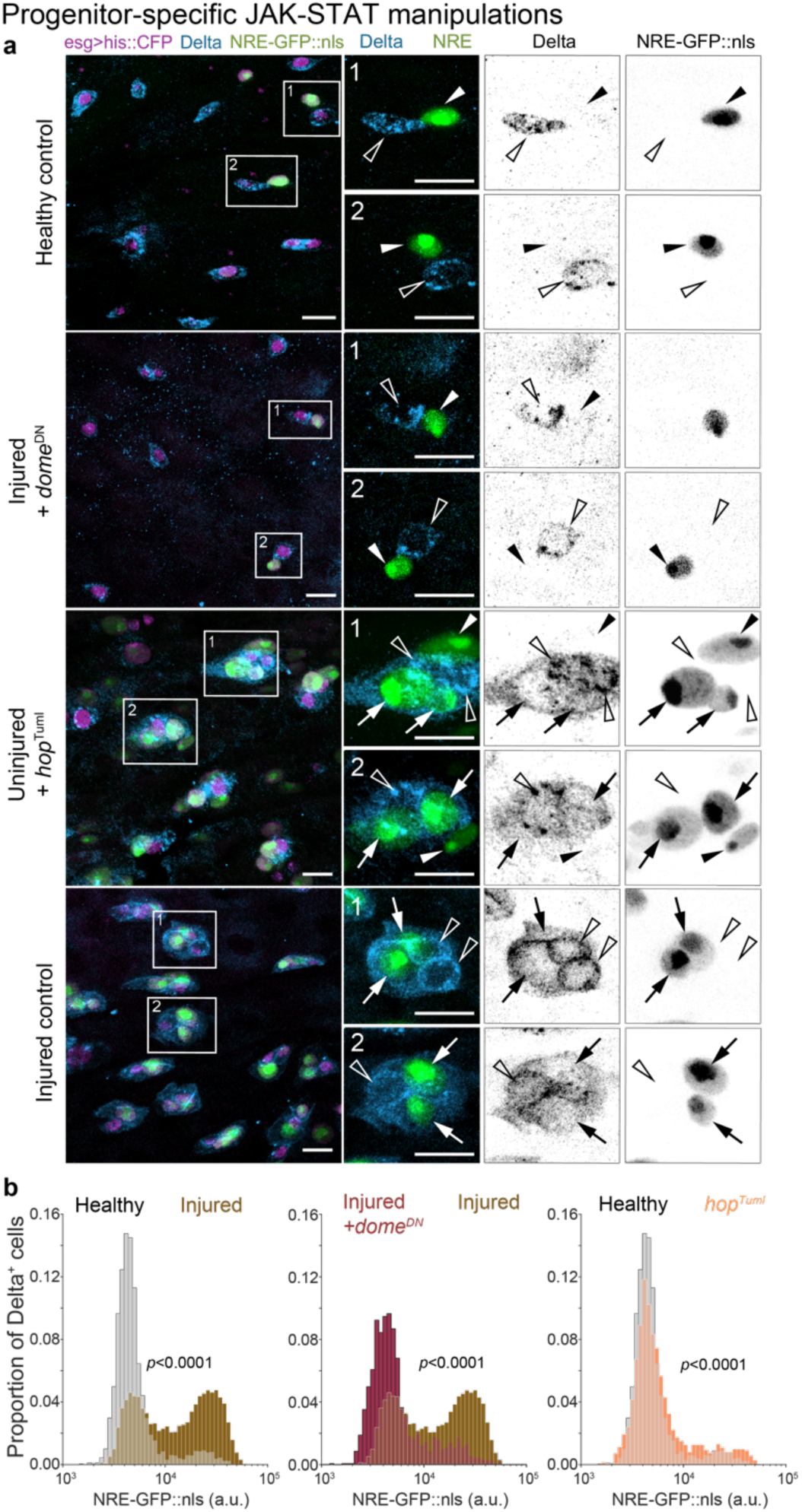
Loss of Notch-Delta feedback occurs through injury-induced activation of Domeless-JAK-STAT. (a) Co-visualization of *NRE-GFP::nls* (green) and Delta immunostain (blue) in progenitor cells (*esg^ts^>his2b::CFP*, magenta) from healthy control guts, injured guts with *dome^DN^*(*esg^ts^>his2b::CFP*, *dome^DN^*), uninjured guts with the activated JAK allele *hop^Tuml^* (*esg^ts^>his2b::CFP*, *hop^Tuml^*), and injured control guts. *dome^DN^*progenitors in injured guts often express either GFP or Delta but not both, similar to healthy control guts. By contrast, *hop^Tuml^*progenitors in otherwise uninjured guts frequently express both GFP and Delta, akin to injured control guts. Arrows indicate Delta^+^, GFP^+^ cells; empty arrowheads indicate Delta^+^, GFP^-^ cells; filled arrowheads indicate Delta^-^, GFP^+^ cells. Scale bars: 10 µm. (b) Notch signaling distributions (*NRE-GFP::nls*) of Delta^+^ progenitors from the same genotypes and conditions in panel a. In injured, *esg^ts^*>*dome^DN^* guts, the proportion of NRE^hi^ cells is reduced compared to injured controls (injured *dome^DN^*– 14%, injured controls - 54%), approaching that of healthy controls (11%). In otherwise uninjured, *esg^ts^>hop^Tuml^*guts, the proportion of NRE^hi^ cells increased relative to healthy controls (uninjured *hop^Tuml^* – 20%, healthy controls – 11%), remaining below injured controls (54%). Healthy control: n=3103 cells; N=19 guts. Injured control: n=6690 cells; N=14 guts. Injured *dome^DN^*: n=1648 cells; N=11 guts. Uninjured *hop^Tuml^*: n=2121 cells; N=11 guts. Samples are from 3 independent experiments. *p*-values from two-sample K-S test. See also Figures S2f, S5e.

We next asked whether JAK-STAT activation alone can disrupt Notch-Delta feedback. Expressing constitutively active JAK (*hopscotch^Tuml^*, hereafter *hop^Tuml^*) caused many progenitors to co-express *NRE-GFP::nls* and Delta, akin to injured controls (Fig. 6a). The proportion of NRE^hi^ cells that were Delta^+^ increased from 16% in healthy controls to 44% in uninjured, *esg^ts^>hop^Tuml^* guts (Figs. S2f, S5e), though remaining below injured controls (69%). Similarly, *hop^Tuml^* increased the proportion of Delta^+^ cells that were NRE^hi^ from 11% to 20%, though again staying below injured controls (54%). These outcomes demonstrate that JAK-STAT activation is sufficient to disrupt Notch-Delta feedback even in the absence of injury.

Together, these findings imply that injury-induced cytokine secretion serves as a tissue-scale damage signal that disrupts progenitor Notch-Delta feedback via Domeless- JAK-STAT activation. Through this relay from damaged cells to differentiating cells, the injured organ couples injury detection to accelerated cell differentiation and regeneration.

## Discussion

Here we uncover an unanticipated mechanism whereby an injured adult epithelial organ hastens the production of differentiated stem cell progeny by exploiting plasticity in the fate-transducing Notch-Delta lateral inhibition circuit (Fig. 7). Our study employs *in vivo* imaging of real-time intestinal repair, single-cell analyses, and dynamical modeling to provide a holistic view of how organs alter fate signaling during injury. While Notch-Delta lateral inhibition is traditionally known as a switch that patterns fate decisions, we find it can also act as a throttle to tune differentiation speed. Cytokines from damaged cells ‘open’ this throttle by eliminating Notch-Delta feedback in stem cell daughters, deactivating the key repressor and prompting sustained Delta expression that accelerates transduction of the fate-determining Notch signal. In this manner, lateral inhibition circuitry elegantly unites temporal and spatial fate control into a single signaling pathway that organs can tune to meet environmental challenges.

**Figure 7:**
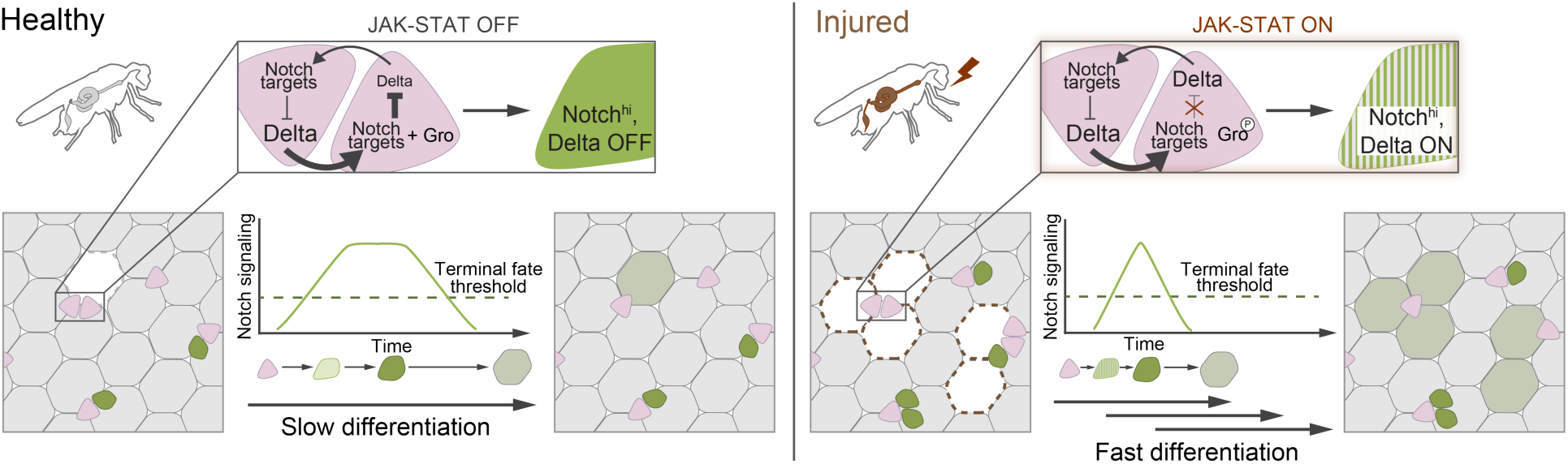
Notch-Delta fate signaling during healthy organ turnover and injury-induced regeneration.

Our finding that cytokine-JAK-STAT is required to eliminate Notch-Delta feedback upends the conventional view that lateral inhibition depends solely on local cell-cell interactions. In the bleomycin-injured fly gut, the primary source of secreted cytokines is damaged enterocytes^2,10,47,49,50^, which do not themselves participate in Notch-Delta circuitry. Through paracrine JAK-STAT activation of progenitor cells, these damaged enterocytes indirectly disrupt Notch-Delta feedback and accelerate cell differentiation. Thus, although lateral inhibition signaling fundamentally involves direct, cell-cell interactions, it is also receptive to long-range signals from third-party cell types.

This mechanism is, to our knowledge, a first demonstration of plasticity in Notch-Delta lateral inhibition circuitry. This form of signaling plasticity differs from prior examples, such as bistability in MAPK cascades^46,79^ and Wnt signaling,^80^ in which signal magnitude is either amplified or abolished. By contrast, lateral inhibition plasticity preserves the overall magnitude of the Notch signal while disrupting its feedback dynamics. This also differs from cell fate plasticity, in which the same molecular signal specifies different fates depending on intensity, duration, or cellular context.^81–88^ Here, both fate outcomes and the level of Notch signal that specifies them remain unchanged (Fig. 2). Instead, this latent plasticity can be invoked for temporal control, providing injured tissues with a parsimonious strategy to rapidly produce mature cells without alternate fate determinants or injury-specific cell types.

We speculate that lateral inhibition plasticity will generalize broadly across biological contexts. Both augmented Notch ligands and faster cell differentiation characterize injury responses in metazoan tissues derived from all three germ layers: ectodermal tissues such as skin^14,15^, endodermal organs such as the intestine^7–11^ and trachea^12,13^, and mesodermal tissues like skeletal and cardiac muscle.^89–91^ Their prevalence raises the possibility that accelerating Notch signaling speed, through either feedback disruption or other mechanisms that increase activating ligands, may be a common regenerative mechanism. Finally, these findings suggest potential therapeutic opportunities to enhance wound repair and organ regeneration through targeted modulation of lateral inhibition plasticity.

## Materials and Methods

### Drosophila husbandry

All experiments were performed on mated adult females. Animals were raised on standard cornmeal–molasses media (water, molasses, cornmeal, agar, yeast, Tegosept, propionic acid). For experiments, we collected adult females post-eclosion and kept them with males in standard cornmeal-molasses vials supplemented with a ∼1cm^2^ sized daub of yeast paste (Red Star, Active Dry Yeast mixed with water) unless otherwise noted.

Genotypes for all fixed experiments included *tub-GAL80^ts^* (i.e., *esg^ts^*>). We reared crosses at 18°C, collected adults on day 0 post-eclosion, then shifted flies to 29°C to inactivate GAL80^ts^ and induce GAL4-mediated expression. Flies were dissected on day 4 post-eclosion.

For live imaging, flies and crosses were kept at 25°C. We collected female flies on day 0 post-eclosion and live-imaged animals on day 3 for all conditions. During all liveimaging experiments, we fed flies via a microcapillary feeder tube with a base recipe of 10% sucrose in water.

### Bleomycin feeding to induce gut injury

To injure the gut, we fed flies Bleomycin (sulfate) (Cayman Chemical #13877).

The injury protocol for live imaging experiments was as follows: Prior to live imaging, flies were fed 25µg/ml bleomycin-laced yeast paste made freshly by mixing dry yeast (Red Star, Active Dry Yeast) with a solution of 25µg/ml bleomycin in water to form a paste. The paste was added to standard cornmeal-molasses vials and refreshed daily for 48 hours prior to live imaging. During live imaging, flies were fed a solution of 10µg/ml bleomycin and 10% sucrose in water via a feeder tube throughout the imaging session.

The injury protocol for all fixed gut experiments except PH3 immunostaining was as follows: 25µg/ml bleomycin-laced yeast paste was prepared as above and provided to flies for 48 hours prior to dissection atop foam plugs wetted with water, refreshed daily. For PH3 staining experiments, 25µg/ml bleomycin-laced yeast paste was administered for 48 hours prior to dissection in standard cornmeal-molasses vials.

### Immunostaining and sample preparation for confocal microscopy

We used the following primary antibodies: rabbit anti-PH3 (EMD Millipore 06-570, 1:400), mouse anti-Delta (DSHB C594-9B – concentrate 1:100, supernatant 1:20). We used the following secondary antibodies: donkey anti-mouse Alexa Fluor 647 (Invitrogen A-31571, 1:400), donkey anti-rabbit Alexa Fluor 555 (Invitrogen A-31572, 1:400). Nuclei were stained with DAPI (Invitrogen D1306, 1:1000 or 1:500). Further details on antibodies and reagents used are provided in Supplementary Table 2.

For PH3 staining (Fig. 2d-f), guts dissected into cold phosphate-buffered saline (PBS) were fixed for 25-30 min at room temperature in 8% formaldehyde (Polysciences 18814-20), 200 mM sodium cacodylate, 100 mM sucrose, 40 mM KOAc, 10 mM NaOAc, and 10 mM EGTA. After fixation, guts were blocked in 0.3% PBT (0.3% Triton X-100 (Sigma-Aldrich X100) in PBS) with 5% normal goat serum (NGS; Capralogics GS0250) for 4 hours at room temperature or overnight at 4°C. Primary and secondary antibodies were incubated in 0.3% PBT + 5% NGS for 4 hours at room temperature or overnight at 4°C. Guts were washed 5 times in PBT between antibody incubations and before mounting.

For staining with mouse anti-Delta (Fig. 2g-k, Fig. 5a-f, Fig. 6) we dissected guts into cold Schneider’s media, fixed in 4% formaldehyde in Schneider’s media at room temperature for 2 hours, and then incubated in 2N HCl in PBS for 20 minutes at room temperature. Next, we washed guts 5x 15 min with Schneider’s media and blocked in 0.3% PBT + 5% NGS at room temperature or overnight at 4°C. We incubated guts in primary antibodies in 0.3% PBT + 5% NGS for 4 hours at room temperature or overnight at 4°C, then washed 5x 15 min in PBS before incubating with secondary antibody. Secondary antibodies were diluted in 0.3% PBT + 5% NGS, and we incubated for 4 hours at room temperature or overnight at 4°C. Finally, we again fixed guts in 4% formaldehyde in PBS for 30 min and washed 4x 15min in PBS before mounting.

We mounted immunostained guts in 3% low-melting 2-hydroxylethyl agarose (Sigma-Aldrich 39346-81-1) and Prolong Gold or Prolong Diamond Antifade mounting media (Thermo Fisher P10144, P36965). We allowed slides to dry covered from light at room temperature for 12-24 hours and stored slides at -20°C until imaging.

### Confocal microscopy

Fixed sample data and images were collected using two microscope systems: (1) a Leica SP8 inverted confocal microscope with a 40x HC PL APO oil objective (Fig. 2a-f,i-k); and (2) a Leica Stellaris8 DIVE confocal microscope with a 20x HC PL APO immersion objective (Fig. 5b,d,f, Fig. 6b) or 40x HC PL APO oil objective (representative figure images: Fig. 2g-h, Fig. 5a,c,e, Fig. 6a). Image acquisition settings were kept consistent within experiments; global shifts in absolute intensity measurements between experiments are due to different microscope systems or different fixation protocols (i.e. Fig. 2d-f PH3 staining vs. Fig. 2i-k Delta staining).

We collected serial optical sections at 2-3µm intervals throughout the entirety of whole-mounted, immunostained guts using Leica Application Suite X (LAS X) (Version 3.5.7.23225). We used Fiji (Version 2.14.0) and Bitplane Imaris x64 (Version 10.1.1) for image analysis.

All image-based quantifications were performed on the R4ab region of the posterior midgut.

### Quantifying *NRE-GFP::nls* activity distributions in fixed tissues

For all *NRE-GFP::nls* intensity measurements, we imaged whole-mounted guts on a Leica SP8 or Stellaris 8 DIVE confocal microscope. Initial .lif files were converted to .ims files and opened in Bitplane Imaris. We used the Add New Surfaces Function in the Surpass Module to generate surfaces for all progenitor nuclei in the *esgGAL4>his2b::CFP* (*esg^+^*) channel. Settings for surface recognition were kept as consistent as possible using the following settings: Smoothing enabled, Surface Grain Size = 0.5µm, Background Subtraction enabled, Diameter of Largest Sphere = 6.00µm, manual threshold value = 4400-max, region growing estimated diameter 3.60µm, ‘Classify Seed Points’ Quality adjusted for each file, ‘Classify Surfaces’ Number of Voxels adjusted for each file 10-∼800 voxels. Surfaces were checked for accuracy and manually edited as needed. For lateral inhibition assay experiments, we identified Delta^+^ cells via immunostaining from the existing *esg^+^* surfaces and processed this Delta^+^, *esg^+^* subset as a separate group. Mean *NRE-GFP::nls* intensity data for both Delta^+^, *esg^+^* and all-*esg^+^* populations was exported as .xlsx and .csv files. Files were loaded in MATLAB (R2024b) and plotted as log-scale histograms with a set bin width interval of 10^0.04^ or 10^0.05^ (Fig. 2a-c). We used the two-sample Kolmogorov-Smirnov (K-S) test to evaluate statistically significant (p<0.05) difference between distributions.

Specifically for measurements of *NRE-GFP::nls* in PH3-stained mitotic cells (Fig. 2d-f), we individually inspected PH3^+^ cells for goodness of fit to the generated surface. Surfaces that overlapped with nuclear signals from neighboring cells were edited to ensure that *NRE-GFP::nls* signal was only coming from the appropriate cell of interest. Cells for which an adjacent, bright GFP^+^ enteroblast interfered with accurate measurement of *NRE-GFP::nls* intensity were excluded from analysis.

### Analyses of NRE-GFP distributions via Gaussian Mixture Model (GMM)

Using the MATLAB fitgmdist() function, we fitted two-component Gaussian mixture models (GMMs) to the distributions of all *esg^+^* progenitor cell *NRE-GFP::nls* intensities for each condition. We took the respective mixing proportions/prior probabilities of the two components to represent the proportions of cells residing in the NRE^low^ and NRE^hi^ peaks (Fig. 2a-c). We took the GMM decision boundary (equal posterior probability threshold) as a proxy for the mean *NRE-GFP::nls* intensity where cells above this threshold are defined as NRE^hi^.

For analysis of PH3^+^ cell *NRE-GFP::nls* distributions (Fig. 2d-f), we again fitted two-component GMMs to the distributions of all *esg^+^* progenitor cell *NRE-GFP::nls* intensities in homeostatic and injured controls, respectively. PH3^+^-cell *NRE-GFP::nls* intensity distributions are displayed as raincloud plots for each condition. We computed the posterior probability prediction of each component (NRE^low^ vs NRE^hi^) for the PH3^+^ datasets against the GMM for their respective condition.

For quantification of progenitor cell Delta-Notch signaling states (Fig. S2), we filtered NRE^hi^ cells from both the all *esg^+^* and the Delta^+^, *esg^+^* datasets, with the latter defined as the Delta^+^, NRE^hi^ group, for each experimental condition using the decision boundary from their respective tissue state GMM (i.e., uninjured background against healthy control GMM, bleomycin-fed against injured control GMM),.

### Single-cell cross-correlation of Notch target and *Delta* mRNAs

We downloaded single-nuclear sequencing 10x Genomics expression matrix files for the *Drosophila* gut from the Fly Cell Atlas^59^ site (https://flycellatlas.org/#data) and parsed them in Python (Version 3.12.3) with Jupyter notebook. Cells from 5do female flies annotated as “intestinal stem cell” and “enteroblast” were parsed out and combined into one all-progenitor pool. We then queried all progenitors for expression levels of *Delta* and the three most highly expressed *E(spl)-C* Notch target genes (-*mα*, -*mβ,* - *m3*)^19,28^ as well as *klumpfuss*, a transcription factor induced specifically in enteroblasts.^92^ Cells with zero levels for both *Delta* and the respective Notch target gene were excluded from further analysis. Normalized expression values were imported into GraphPad Prism 10 (Version 10.3.1) for plotting and correlation analysis.

### Modeling Notch-Delta lateral inhibition

We considered that the active Notch level of a cell is an increasing function of the Delta level of neighboring cells, and that the Delta level of a cell is a decreasing function of the active Notch level of that cell. We formulate this interaction between pairs of cells using standard mathematical models of Notch-Delta lateral inhibition.^27,34^ In its dimensionless form, the equations can be written as:

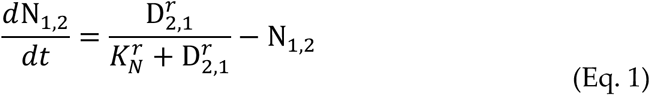

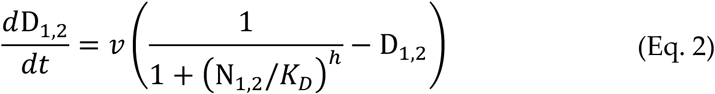

where the subscript denotes the Notch/Delta of cell 1 or 2. In these equations, K_N_ is the dimensionless threshold of Notch activation by Delta ligand of neighboring cell, and K_D_ is the dimensionless threshold of Delta inhibition by activated Notch of the same cell. The parameter 𝑣 is the ratio of degradation rate of Notch to Delta, which following previous work, we are assuming is equal to one.^27,34,63^ According to Guisoni et al., 2017,^27^ K_N_ is inversely related to the contact area between two cells. More generally, K_N_ dictates the intercellular aspect of Notch-Delta interaction, while K_D_ dictates the intracellular aspect. The parameters r and h are the hill coefficients for Notch activation and Delta inhibition and are considered r=h=2 to account for the cooperative nature of these processes.^34^

To simulate the activation of a downstream Notch reporter, we assumed that reporter expression is directly related to activated Notch levels:

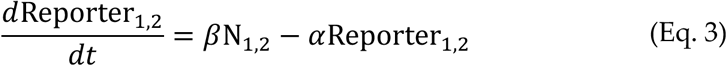

Where *β* is the maximal production rate of reporter, and *α* is the degradation rate of reporter. Since the dimensionless Notch levels range between zero and one, the above equation would show no reporter expression prior to Notch activation and the reporter levels would reach steady state at *β*/*α* after full Notch activation. Immediately after Notch activation, the reporter expression is dominated by production rate and invariable to the degradation rate. Therefore, we approximate the reporter level by the following, with *β* = 1:

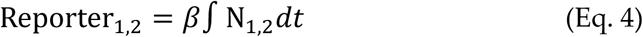

### Modeling simulation conditions

We numerically solved the above equations to derive the time dynamics of Notch and Delta using the odeint function from python’s scipy library. Cells are initially considered to be low Notch and high Delta. To break the symmetry between the two cells, cell 2 has a slightly higher initial Notch level than cell 1 (0.010 versus 0.011). We used a plausible range of K_N_ and K_D_ parameters to study the behavior of Notch-Delta dynamics.^27,63,93,94^ Particularly, data fitted to wildtype cells from Guisoni et al., 2017^27^ Figure 4 shows a K_D_ range of 0.2-0.3, and a K_N_ range of 0.1-10.

### Windowmount live imaging

We performed Windowmount live imaging of the *Drosophila* midgut as previously described.^44^ Briefly, we glued female flies to the imaging apparatus and opened a window in the dorsal cuticle of the abdomen. The R4 region of the midgut was identified, nudged through the cuticular window, and stabilized with 3% agarose before being bathed with live imaging media (recipe below). We then imaged the exposed region of the midgut using an upright Leica SP5 multi-photon confocal microscope with a 20x water immersion objective (Leica HCX APO L 20x NA 1.0). We fed flies 10% sucrose either with or without 25 µg/ml bleomycin through a microcapillary feeder tube during the entire imaging session. Movies were captured at room temperature (20–25°C). Confocal stacks were acquired with a Z-step of 2.98 µm at 7.5min intervals and typically contained ∼35-40 slices.

### Live imaging media recipe

All guts analyzed of NRE-GAL4>TransTimer genotypes used the following recipe adapted from Marco Marchetti and Bruce Edgar (University of Utah), who have since published an updated version^95^: 61.5mM L-Glutamic acid monosodium salt (made in Schneider’s media), 55.5mM Trehalose (made in Schneider’s media), 2.2mM N-Acetyl Cysteine (made in water), 1.1mM Tri-sodium Citrate (made in Schneider’s media), 11% Fetal Calf Serum (or fetal bovine serum (FBS)), Schneider’s media, Penicillin-streptomycin 0.55%. Stocks of the above ingredients were made in advance, filter sterilized using a 0.2µm syringe filter, and stored at 4°C for up to 3 months. We made live imaging media fresh on the day of imaging. Media was stored at 4°C and used until the next day if needed.

### Live imaging movie registration

After acquisition, movies were processed on a Windows computer (Windows 10 Education) with a 3.70 GHz quad-core Intel Xeon processor and 128 GB memory. LIF files (*.lif) from Leica Application Suite: Advanced Fluorescence were uploaded into Fiji as a hyperstack for registration. To correct for X-Y drift, movies were converted to RGB files and processed with the Fiji plugin StackReg.^96^ To correct for global volume movements, movies were processed with the Fiji plugin Correct 3D Drift.^97^ We evaluated movies for viability based on criteria established in Martin et al., 2018.^44^

### Live imaging analyses: Cell identification, tracking, and TransTimer quantification in Imaris

To perform cell tracking, processed and registered movies were converted from .tiff format to .ims file format using the Bitplane Imaris File Converter software. We performed cell segmentation in Bitplane Imaris 9.2.0 using the TransTimerRFP channel to generate 3D “spots” with the “Spots” module. All spots were generated using a standardized spot diameter of 9.02 mm. We used the Brownian motion tracking algorithm to track cell surfaces and spots for all labeled cells across all movie time points. Any errors in cell surface generation and tracking were visually inspected and corrected. Once cell recognition was verified for all cells for all time points, we exported individual cell measurements for mean intensity GFP and mean intensity RFP as Microsoft Excel files. For each channel within a movie, mean intensity values were normalized to a 0-to-1 scale by setting the maximum intensity measurement to 1. Data was imported into MATLAB or GraphPad Prism for analysis.

We note that, from a technical standpoint, these real-time measurements of TransTimer dynamics provide useful insight for interpreting TransTimer data in fixed samples.

### Calculating rate of NRE>TransTimerGFP signal change

After we standardized normalizing TransTimerGFP values over time for each movie, we plotted tracks over time for each cell and smoothed the data using the ‘rlowess’ method and a moving time-average spanning 5 timepoints in MATLAB. Cells were excluded from further analysis if the average of the first half of the data points in the track were <0.1 mean GFP intensity. Cells that still had visible TransTimerRFP expression but had TransTimerGFP intensity < 0.1 were designated as recently Notch-OFF cells that were excluded from slope analysis. Cells were excluded from further analysis if: (1) fewer than 8 data points were collected or (2) noise in raw data measurements precluded meaningful analysis. To enable accurate slope analysis of tracks with distinct positive and negative slope segments, we split tracks into two parts at the maximum value of the smoothed data. Data before the maximum should have a positive slope, and after, a negative slope. We then fit the equation (y=mx+b) to the smoothed data. Slope measurements were separated into positive and negative slopes for plotting and comparison.

### Statistical analyses

Statistical analyses and histogram plotting for fixed *NRE-GFP::nls* quantifications were done in MATLAB and edited in Adobe Illustrator (Version 29.0). For comparisons of *NRE-GFP::nls* distributions, we used the two-sample Kolmogorov-Smirnoff (K-S) test to assess statistical significance.

All plots for TransTimer tracks and slopes (Fig. 4f-l), single-cell cross correlation plots (Fig. S3), and violin plots of Delta^+^, NRE^hi^ proportions (Fig. S5) were made in GraphPad Prism 10 and edited in Adobe Illustrator. For comparisons of distributions of cell slopes, we used unpaired two-tailed Mann-Whitney tests to assess median and statistical significance. For comparisons of cell numbers, we used unpaired Student’s twotailed *t*-tests to assess mean and statistical significance. For single-cell cross-correlation (Fig. S3), we used Pearson correlation coefficients (*r*) and *p*-values (two-tailed *t*-test) to assess correlation and statistical significance. For Delta^+^, NRE^hi^ violin plots (Fig. S5), we used ordinary one-way ANOVA with post-hoc Tukey’s multiple comparisons test to assess mean and statistical significance.

The number of experimental replicates for each assay is indicated in the figure legends. Statistical tests used are indicated in the figure legends.

For all experiments, randomization was not relevant/not performed. Data collection and analysis were not performed blind to the conditions of the experiments. All data were acquired and processed identically and in parallel. We used GraphPad Prism 10 (Version 10.3.1), Microsoft Excel 365 (Version 16.90), MATLAB (R2024b), and Python (Version 3.12.3) for statistics and graph generation. We used Adobe Illustrator (Version 29.0) for figure assembly.

## Supporting information

Modeling Supplement

## Data availability

All data that support the findings of this study are available from the authors upon reasonable request.

## Code availability

Modeling code is publicly available on GitHub: https://github.com/htmsun/Sun_Modeling_Code. Other figure-assembly scripts are available from the authors upon reasonable request.

## Acknowledgements and funding

We are grateful to J. de Navascues, N. Perrimon, D. Bilder, D. Montell, S.X. Hou, and *Drosophila* stock centers (Bloomington *Drosophila* Stock Center (NIH P40OD018537), Vienna *Drosophila* Resource Center^98^ for fly stocks, B.A. Edgar for sharing reagent information, and J. Mulholland and K. Lee for microscopy support. Confocal microscopy was performed at the Stanford Beckman Cell Sciences Imaging Facility (RRID:SCR_017787: NIH 1S10OD032300-01, NIH 1S10OD010580-01A1). We thank A. Jacobo, J.E. Ferrell, J. de Navascues, and B. Wang for invaluable discussions and A. Jacobo, Z. Scott, E.G. Magny, J.D. Axelrod, and J.M. Knapp for comments on the manuscript.

H.-T. S. was supported by a NIH T32 (T32GM007790) and an American Heart Association Predoctoral Fellowship (23PRE1012896). E.N.S was supported by NIH T32GM007790 and a National Science Foundation Graduate Research Fellowship (NSF GRFP, DGE-1656518). S.T. was supported by an NSF GRFP (DGE-2146755) and a Stanford Graduate Fellowship. C.F.L. was supported by a Stanford Deans’ Postdoctoral Fellowship. L.H. was supported by the Damon Runyon Cancer Research Foundation and NIH R21DA039582 to N. Perrimon. M.M. was supported by NIH R01GM124434 and NIH R35GM140900 to B.A. Edgar. S.X. was supported by a NIH K99 (1K99GM138712). This work was supported by NIH R01GM116000-01A1, NIH R35GM141885-01, and NIH R01DK128485-01A1 to L.E.O. L.E.O. is an investigator of the Chan-Zuckerberg Biohub—San Francisco.

## Contributions

E.N.S and L.E.O. conceived and designed the initial study. H.-T.S. and L.E.O. conceived and designed the current study. H.-T.S. and S. G. van D. performed and analyzed the fixed tissue experiments in this study. S.T. performed the modeling experiments in this study. S.T. and C.F.L. analyzed the modeling experiments in this study. E.N.S. and Y.-H.S. performed the live imaging experiments in this study. E.N.S. and A.L. analyzed the live imaging experiments in this study. E.N.S., A.L. and J.I. processed the live imaging movies in this study. L.H. generated the TransTimer line for live imaging experiments in this study. M.M. formulated the recipe for live imaging media. H.-T.S., E.N.S., and L.E.O. prepared the figures. L.E.O., H.-T. S., S.T., and C.F.L. wrote the manuscript. L.E.O., H.-T. S, C.F.L., E.N.S., and S.X. revised the manuscript. L.E.O. supervised the project.

## Supplemental Figures + Captions, and Movie Captions

In order of appearance in the manuscript

**Supplemental Figure 1:**
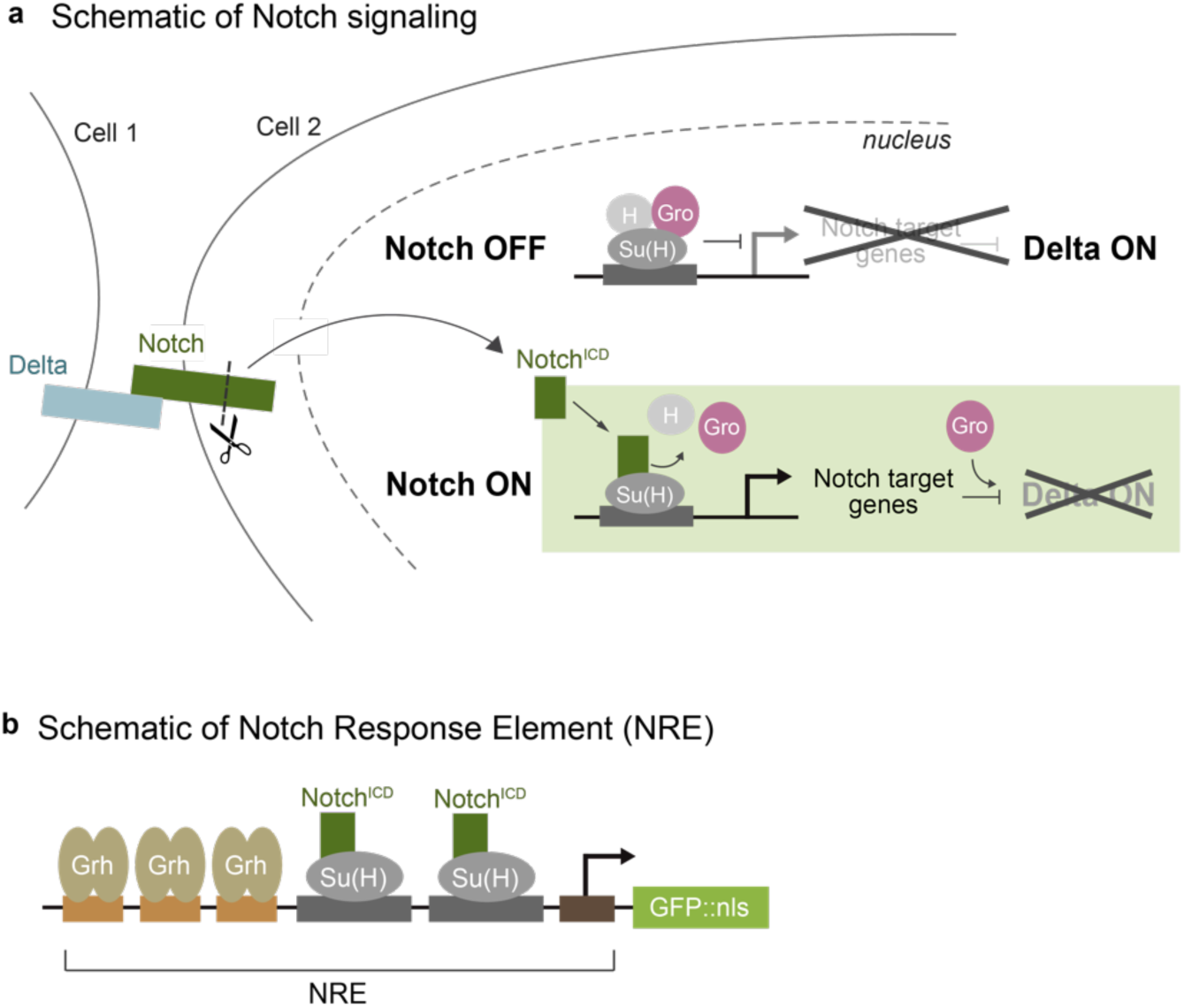
Molecular regulation of Notch target genes and the Notch Response Element (NRE). (a) Simplified schematic of Notch target regulation. In the inactive state (Notch OFF), Suppressor of Hairless (Su(H)) bound to DNA sites (gray boxes) recruits co-repressors Hairless (H) and Groucho (Gro), silencing Notch targets while permitting Delta expression. In the active state, Delta ligand (blue) binds Notch receptor (green) (Notch ON), releasing the Notch intracellular domain (Notch^ICD^). Notch^ICD^ enters the nucleus, binds Su(H), and displaces H/Gro. The Notch^ICD^/Su(H) complex then drives Notch target gene expression. Notch targets, together with Gro, repress Delta transcription. (b) Structure of the Notch Response Element (NRE) reporter. Sensitive detection of Notch activation is conferred by the combination of two Su(H) binding sites with three transcriptional activator Grainyhead (Grh) binding sites (GBE).^45^ The NRE drives expression of nuclear GFP (GFP::nls) in all figures except Figure 4, where it drives GAL4.

**Supplemental Figure 2:**
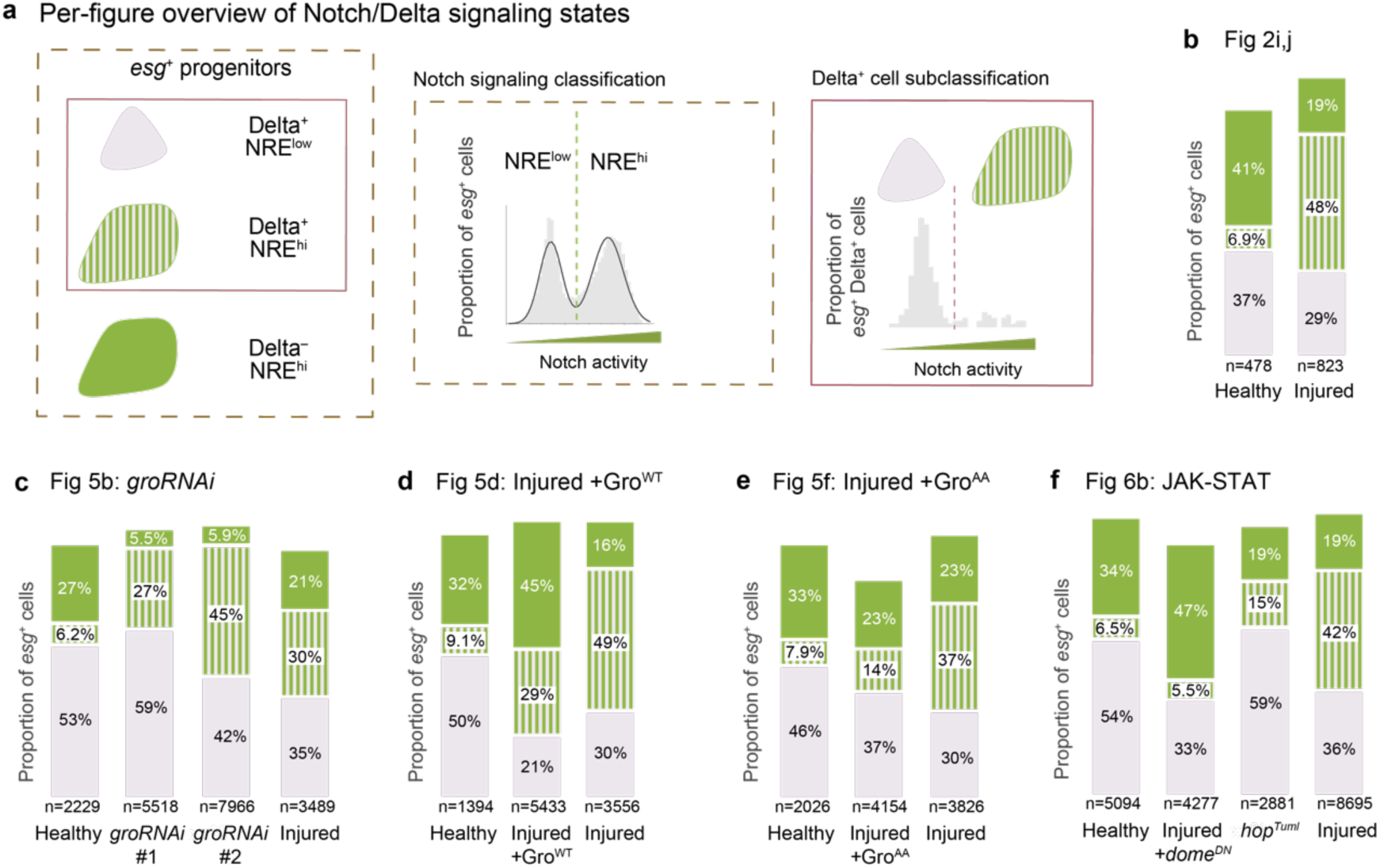
Overview of Notch/Delta signaling states across experimental conditions. (a) Classification framework for Notch/Delta signaling states in midgut progenitors (*esg^+^*). (b-f) Quantitation of signaling states (percent of total *esg^+^*cells) for: (b) Fig 2i,j: Healthy vs injury; (c) Fig 5b: *gro* RNAi (uninjured); (d) Fig 5d: Injury + *gro^WT^*; (e) Fig 5f: Injury + *gro^AA^*; (f) Fig 6b: JAK-STAT pathway perturbations. Independent healthy and injured control guts were included with each experiment. Values shown as percentage of total *esg^+^* cells. Delta^−^, NRE^low^ cells excluded as they do not signal, so proportions sum to <100%.

**Supplemental Figure 3:**
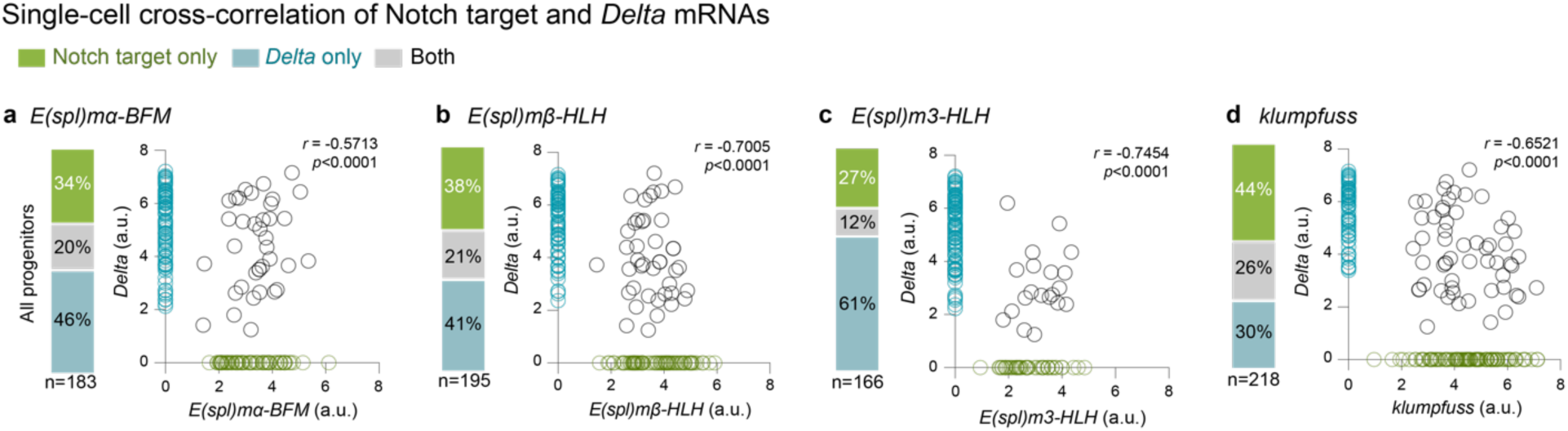
Anti-correlation of Delta and Notch-target mRNAs in healthy-gut progenitors. Single-cell expression analyses of *Delta* and four major midgut Notch target genes. We queried progenitors from the Fly Cell Atlas^59^ for expression levels of *Delta* and the three most highly expressed *E(spl)-C* Notch target genes (-*mα*, -*mβ,* -*m3*)^19,28^ as well as *klumpfuss*, a transcription factor induced specifically in enteroblasts.^92^ (a) *E(spl)mα-BFM*, (b) *E(spl)mβ-HLH*, (c) *E(spl)m3-HLH*, and (d) *klumpfuss*. Stacked bars quantify proportions of progenitor cells that express only *Delta* (blue), only Notch target (green), or both (gray). Scatter plots show *Delta* versus Notch target mRNA levels per cell, with corresponding color-coding. Data from 5-day-old, mated female flies.^59^ See Methods. *r* = Pearson’s correlation coefficient; *p*-values from two-tailed t-test.

**Supplemental Figure 4:**
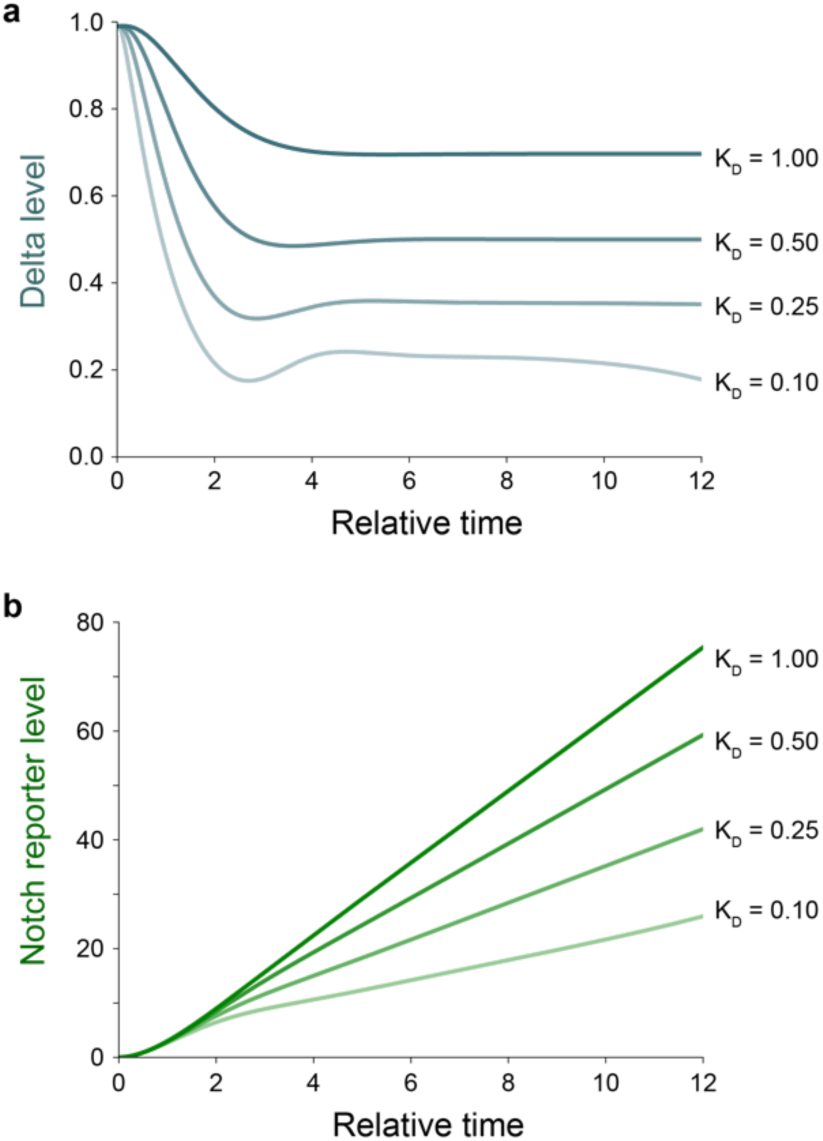
Delta and Notch signaling dynamics across K_D_ values. Simulated time evolution of (a) Delta levels and (b) Notch reporter levels at the indicated K_D_ values. Increased K_D_ produces higher levels of both Delta and Notch reporter. K_N_ =0.5 in all simulations.

**Movie 1: 20.5-hour live imaging movie of a healthy NRE>TransTimer gut**

See Figure 4d. Two-channel, wide-field, volumetric movie of a healthy NRE>TransTimer gut. White lines initially outline the gut boundaries. NRE>TransTimerGFP (green) marks cells with active Notch signaling. NRE>TransTimerRFP (magenta) marks recent Notch signaling activity. Scale bar, 50µm.

**Movie 2: 20.5-hour live imaging movie of an injured NRE>TransTimer gut**

See Figure 4e. Two-channel, wide-field, volumetric movie of an injured NRE>TransTimer gut. White lines initially outline the gut boundaries. NRE>TransTimerGFP (green) marks cells with active Notch signaling. NRE>TransTimerRFP (magenta) marks recent Notch signaling activity. Scale bar, 50µm.

**Movie 3: Healthy NRE>TransTimer cell exhibiting NRE upregulation**

See Figure 4f, Cell 1. Cell in frame increases both NRE>TransTimerGFP (first panel, green; second panel, inverted gray) and NRE>TransTimerRFP (first panel, magenta; third panel, inverted gray) signal over the course of the 20.5-hour movie. Each time point is the projection of a confocal z-stack. Scale bar, 5µm.

**Movie 4: Healthy NRE>TransTimer cell exhibiting sustained NRE signal**

See Figure 4g, Cell 2. The centermost GFP+ cell in frame exhibits sustained NRE>TransTimerGFP (first panel, green; second panel, inverted gray) and NRE>TransTimerRFP (first panel, magenta; third panel, inverted gray) signal over the course of the 20.5-hour movie. Each time point is the projection of a confocal z-stack. Scale bar, 5µm.

**Movie 5: Healthy NRE>TransTimer cell exhibiting NRE downregulation.**

See Figure 4h, Cell 3. The centermost GFP+ cell in frame (denoted by white arrow) exhibits decreasing NRE>TransTimerGFP (first panel, green; second panel, inverted gray) and NRE>TransTimerRFP (first panel, magenta; third panel, inverted gray) signal over the course of the 20.5-hour movie. Each time point is the projection of a confocal z-stack. Scale bar, 5µm.

**Movie 6: Injured NRE>TransTimer cell exhibiting both NRE upregulation and downregulation.**

See Figure 4i, Cell 4. Cell in frame exhibits both increasing and decreasing NRE>TransTimerGFP (first panel, green; second panel, inverted gray) and NRE>TransTimerRFP (first panel, magenta; third panel, inverted gray) signal in the course of the 20.5-hour movie. Each time point is the projection of a confocal z-stack. Scale bar, 5µm.

**Supplemental Figure 5:**
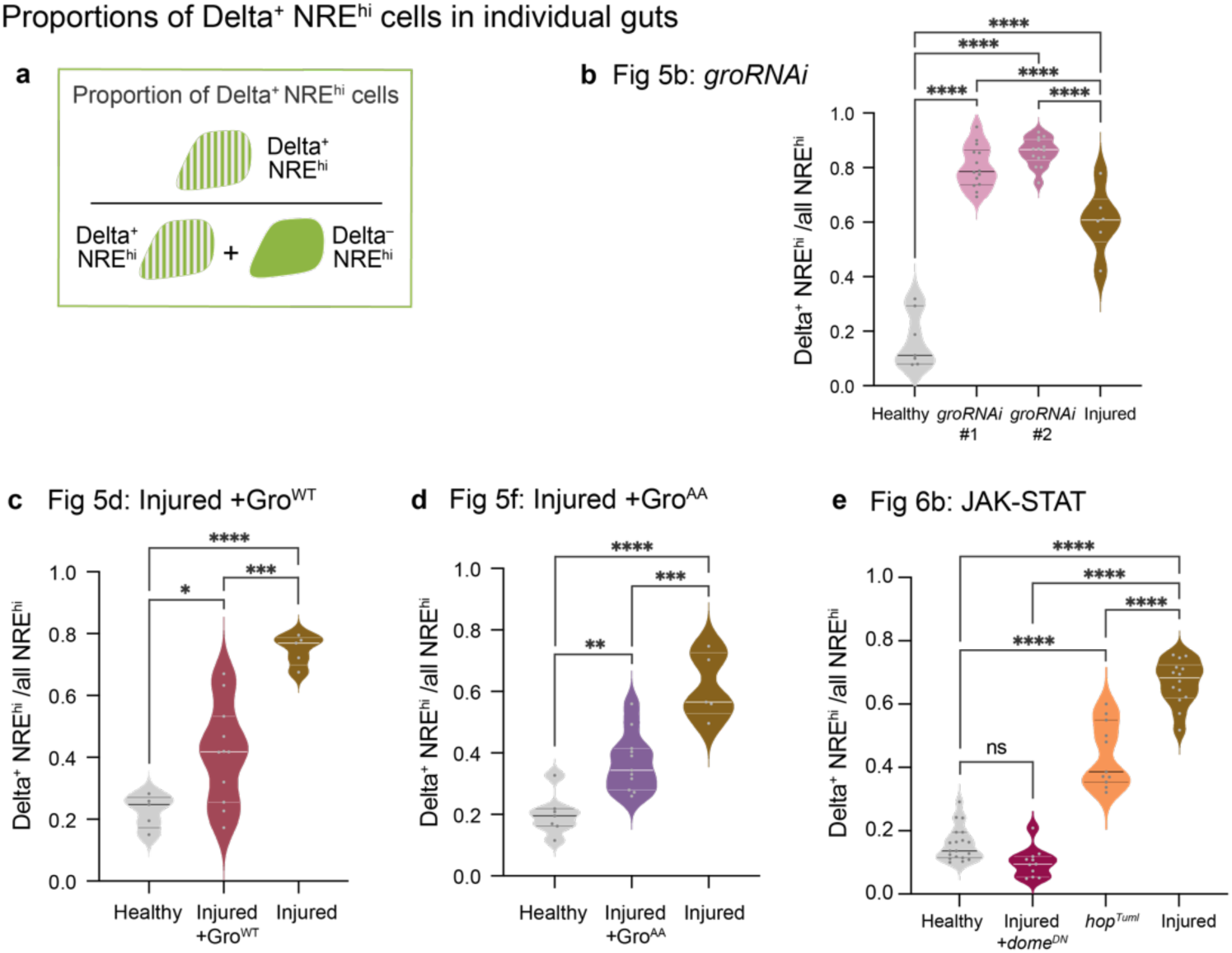
Analysis of the proportion of Delta^+^, NRE^hi^ enteroblasts on a per-gut basis across conditions. (a) Schematic of calculation for proportion of NRE^hi^ cells that are Delta^+^. Violin plots of the proportion of NRE^hi^ cells that are Delta^+^ for data corresponding to: (b) Fig 5b, (c) Fig 5d, (d) Fig 5f, and (e) Fig 6b. Each dot represents one gut. Horizontal lines represent median and 25th, 75th percentiles. *p-*values, one-way ANOVA with *post hoc* Tukey test for multiple comparisons. ns, not significant; *, *p*<0.05; **, *p*<0.01; ***, *p*<0.001; ****, *p*<0.0001.

**Table 1.** Genotypes in Figure Panels.

| FIGURE | GENOTYPE |
| --- | --- |
| Fig 1c | esgGAL4, UAS-his2b::CFP, GBE-Su(H)-GFP::nls/+; ubi-E-cadherin::YFP/+ |
| Fig 2a-k | w<sup>1118</sup>/+; esgGAL4, UAS-his2b::CFP, GBE-Su(H)-GFP::nls/+; tubGAL80<sup>ts</sup>/+ |
| Fig 4d-l | NRE>TransTimer: GBE-Su(H)GAL4/CyO; UAS-TransTimer/TM3 |
| Fig 5a,b | w<sup>1118</sup>/+; esgGAL4, UAS-his2b::CFP, GBE-Su(H)-GFP::nls/+; tubGAL80<sup>ts</sup>/+,<br>esgGAL4, UAS-his2b::CFP, GBE-Su(H)-GFP::nls/+; tubGAL80<sup>ts</sup>/UAS-groRNAi #1,<br>esgGAL4, UAS-his2b::CFP, GBE-Su(H)-GFP::nls/UAS-groRNAi #2; tubGAL80<sup>ts</sup>/+ |
| Fig 5c,d | esgGAL4, UAS-his2b::CFP, GBE-Su(H)-GFP::nls/UAS-Gro<sup>WT</sup>; tubGAL80<sup>ts</sup>/+ |
| Fig 5e,f | w<sup>*</sup>/+; esgGAL4, UAS-his2b::CFP, GBE-Su(H)-GFP::nls/UAS-Gro<sup>AA</sup>; tubGAL80<sup>ts</sup>/+ |
| Fig 6a,b | w<sup>1118</sup>; esgGAL4, UAS-his2b::CFP, GBE-Su(H)-GFP::nls/+; tubGAL80<sup>ts</sup>/+,<br>w<sup>*</sup>/+; esgGAL4, UAS-his2b::CFP, GBE-Su(H)-GFP::nls/UAS-dome<sup>DN</sup>; tubGAL80<sup>ts</sup>/Dr<br>UAS-hop<sup>TumI</sup>/+; esgGAL4, UAS-his2b::CFP, GBE-Su(H)-GFP::nls/+; tubGAL80<sup>ts</sup>/+, |

**Table 2.**
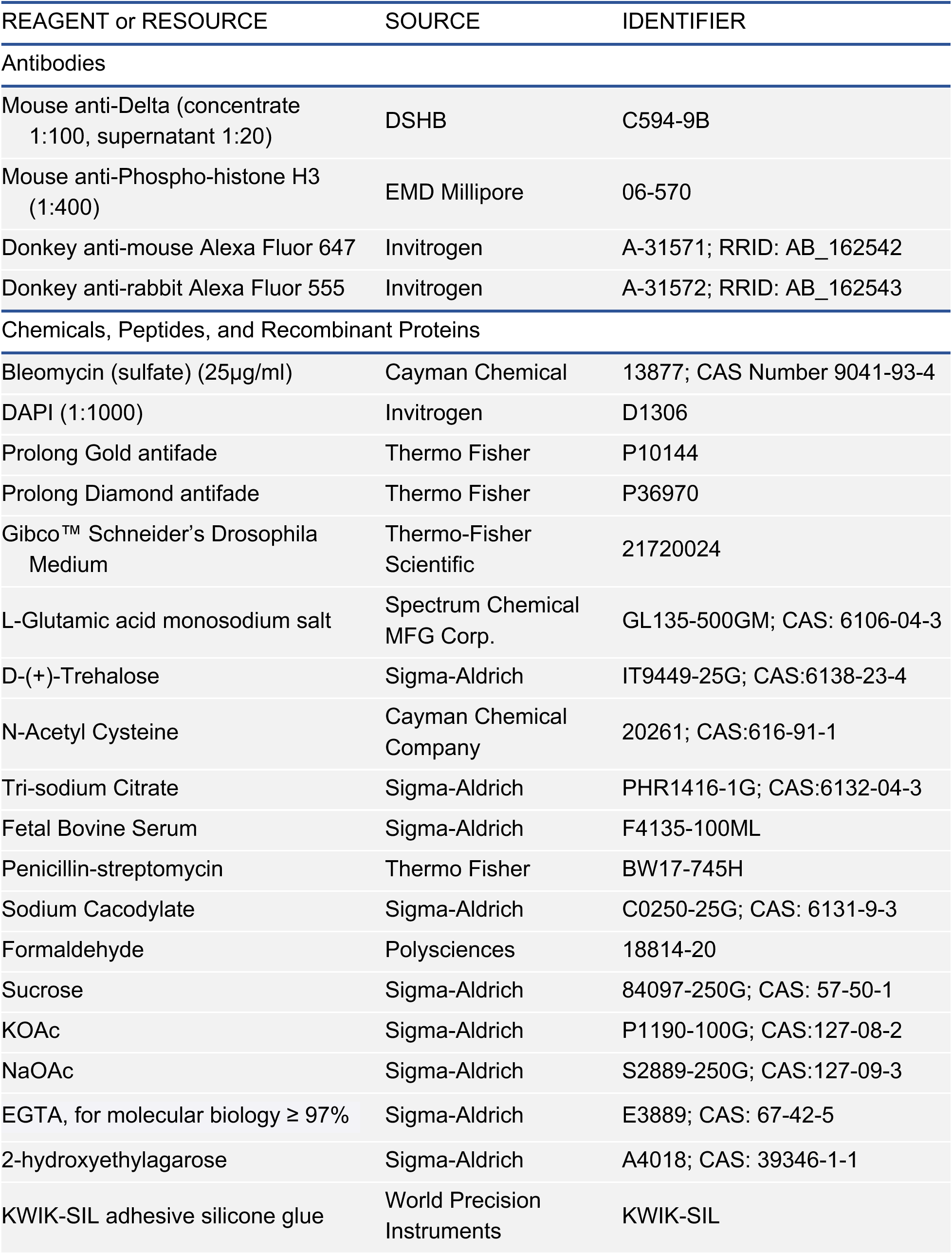
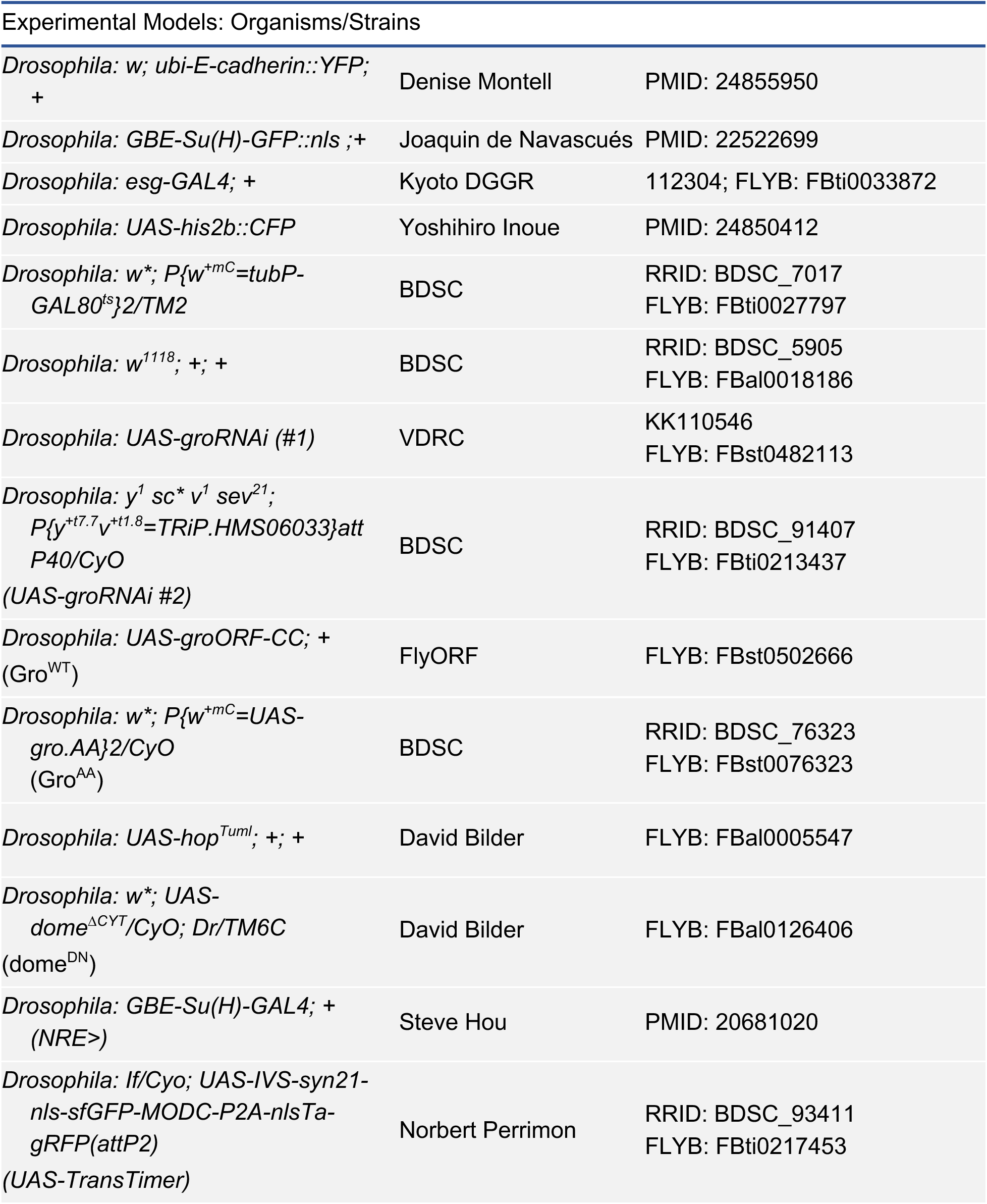
Reagents and Resources.

| REAGENT or RESOURCE | SOURCE | IDENTIFIER |
| --- | --- | --- |
| **Antibodies** |  |  |
| Mouse anti-Delta (concentrate 1:100, supernatant 1:20) | DSHB | C594-9B |
| Mouse anti-Phospho-histone H3 (1:400) | EMD Millipore | 06-570 |
| Donkey anti-mouse Alexa Fluor 647 | Invitrogen | A-31571; RRID: AB_162542 |
| Donkey anti-rabbit Alexa Fluor 555 | Invitrogen | A-31572; RRID: AB_162543 |
| **Chemicals, Peptides, and Recombinant Proteins** |  |  |
| Bleomycin (sulfate) (25µg/ml) | Cayman Chemical | 13877; CAS Number 9041-93-4 |
| DAPI (1:1000) | Invitrogen | D1306 |
| Prolong Gold antifade | Thermo Fisher | P10144 |
| Prolong Diamond antifade | Thermo Fisher | P36970 |
| Gibco™ Schneider's Drosophila Medium | Thermo-Fisher Scientific | 21720024 |
| L-Glutamic acid monosodium salt | Spectrum Chemical MFG Corp. | GL135-500GM; CAS: 6106-04-3 |
| D-(+)-Trehalose | Sigma-Aldrich | IT9449-25G; CAS:6138-23-4 |
| N-Acetyl Cysteine | Cayman Chemical Company | 20261; CAS:616-91-1 |
| Tri-sodium Citrate | Sigma-Aldrich | PHR1416-1G; CAS:6132-04-3 |
| Fetal Bovine Serum | Sigma-Aldrich | F4135-100ML |
| Penicillin-streptomycin | Thermo Fisher | BW17-745H |
| Sodium Cacodylate | Sigma-Aldrich | C0250-25G; CAS: 6131-9-3 |
| Formaldehyde | Polysciences | 18814-20 |
| Sucrose | Sigma-Aldrich | 84097-250G; CAS: 57-50-1 |
| KOAc | Sigma-Aldrich | P1190-100G; CAS:127-08-2 |
| NaOAc | Sigma-Aldrich | S2889-250G; CAS:127-09-3 |
| EGTA, for molecular biology ≥ 97% | Sigma-Aldrich | E3889; CAS: 67-42-5 |
| 2-hydroxyethylagarose | Sigma-Aldrich | A4018; CAS: 39346-1-1 |
| KWIK-SIL adhesive silicone glue | World Precision Instruments | KWIK-SIL |
| Experimental Models: Organisms/Strains |  |  |
| *Drosophila*: *w*; *ubi-E-cadherin::YFP*;<br>+ | Denise Montell | PMID: 24855950 |
| *Drosophila*: *GBE-Su(H)-GFP::nls*; + | Joaquin de Navascués | PMID: 22522699 |
| *Drosophila*: *esg-GAL4*; + | Kyoto DGGR | 112304; FLYB: FBti0033872 |
| *Drosophila*: *UAS-his2b::CFP* | Yoshihiro Inoue | PMID: 24850412 |
| *Drosophila*: *w\**; *P{w<sup>+mC</sup>=tubP-GAL80<sup>ts</sup>}2/TM2* | BDSC | RRID: BDSC_7017<br>FLYB: FBti0027797 |
| *Drosophila*: *w<sup>1118</sup>*; +; + | BDSC | RRID: BDSC_5905<br>FLYB: FBal0018186 |
| *Drosophila*: *UAS-groRNAi* (#1) | VDRC | KK110546<br>FLYB: FBst0482113 |
| *Drosophila*: *y<sup>1</sup> sc\* v<sup>1</sup> sev<sup>21</sup>*;<br>*P{y<sup>+t7.7</sup> v<sup>+t1.8</sup>=TRiP.HMS06033}att*<br>*P40/CyO*<br>(*UAS-groRNAi* #2) | BDSC | RRID: BDSC_91407<br>FLYB: FBti0213437 |
| *Drosophila*: *UAS-groORF-CC*; +<br>(*Gro<sup>WT</sup>*) | FlyORF | FLYB: FBst0502666 |
| *Drosophila*: *w\**; *P{w<sup>+mC</sup>=UAS-gro.AA}2/CyO*<br>(*Gro<sup>AA</sup>*) | BDSC | RRID: BDSC_76323<br>FLYB: FBst0076323 |
| *Drosophila*: *UAS-hop<sup>Tuml</sup>*; +; + | David Bilder | FLYB: FBal0005547 |
| *Drosophila*: *w\**; *UAS-dome<sup>ΔCYT</sup>/CyO*; *Dr/TM6C*<br>(*dome<sup>DN</sup>*) | David Bilder | FLYB: FBal0126406 |
| *Drosophila*: *GBE-Su(H)-GAL4*; +<br>(*NRE>*) | Steve Hou | PMID: 20681020 |
| *Drosophila*: *lf/Cyo*; *UAS-IVS-syn21-nls-sfGFP-MODC-P2A-nlsTagRFP(attP2)*<br>(*UAS-TransTimer*) | Norbert Perrimon | RRID: BDSC_93411<br>FLYB: FBti0217453 |

