## Supplementary material for "Organ injury accelerates stem cell differentiation by modulating a fate-transducing lateral inhibition circuit": Modeling Supplement

### Three-cell Model of Notch-Delta Lateral Inhibition

#### 1 Introduction

Upon injury, Drosophila intestinal cells exhibit a unique Delta-positive Notch-positive population. Previous work using a two-cell model of Notch Delta signaling adapted from Guisoni et al. 2017 was sufficient to recreate this observation. More specifically, disruption of the inhibitory loop between Notch and Delta was necessary to create this effect, and furthermore, this perturbation is predicted to increase Notch activation speed, which was experimentally proven to be true.

It was also observed that upon injury, there is an increased instance of multiple cells contacting each other, suggesting that the two-cell model is no longer accurate. Additionally, it was suggested that the emergence of the Delta-positive Notch-positive population may be due to this clustering effect, and not by the disruption of Delta inhibition as seen in the two-cell model. To address these concerns, we have generated a three-cell model of Notch Delta signaling to explore whether the addition of another cell to the model is sufficient in generating a high-Delta high-Notch state.

#### 2 Theoretical Framework

To generate a three-cell model, we decided to extend the two-cell model. Briefly, the two-cell model from Guisoni et al. 2017 in its reduced form has the following equations:

$$\frac{d\mathrm{N}_{1,2}}{dt} = \frac{\mathrm{D}_{2,1}^r}{K_N^r + \mathrm{D}_{2,1}^r} - \mathrm{N}_{1,2} \tag{1}$$

$$\frac{d\mathrm{D}_{1,2}}{dt} = v\left(\frac{1}{1 + \left(\mathrm{N}_{1,2}/K_D\right)^h} - \mathrm{D}_{1,2}\right) \tag{2}$$

Where $v$ is the ratio of Notch and Delta degradation rates and generally assumed to equal to one. The main parameters in this model are $K_N$ and $K_D$ which are respectively the threshold of Notch activation by neighboring Delta, and Delta inhibition by Notch of the same cell. According to Guisoni et al. 2017, $K_N$ is inversely related to cell-cell contact area, in other words, cells with high contact area have more Notch receptors activated by neighboring Delta, and thus lower $K_N$. Conversely, cells with little contact area do not have much Notch activation, and thus higher $K_N$.

To extend the Guisoni et al. model to three cells, we need to change the Hill function that describes Notch activation in equation 1. If cell 2 is contacting cell 1, it should be sufficient to activate cell 1 independent of cell 3. If cell 2 is activating few receptors independently and cell 3 is activating few receptors, this effect should be additive. This means that if cells 2 and 3 have the same number of Delta ligands, their combined effect would be the same as cell 2 making twice the amount of contact alone. In other words, Notch activation of cell 1 should follow an OR logic, where activation from either cell 2 or 3 is sufficient, and the combined effect should also be additive.

To employ this OR logic, we take inspiration from Guisoni et al. 2017's definition of $K_N = c/A$, where c is a constant and A is the area of contact. By plugging this definition back into equation (1), we get the following activation function formulation:

$$\frac{(A \cdot D)^r}{c^r + (A \cdot D)^r} \tag{3}$$

This equation can be rewritten to account for two pairs of interactions:

$$\frac{(A_{1-2}D_2+A_{1-3}D_3)^r}{c^r+(A_{1-2}D_2+A_{1-3}D_3)^r} \tag{4}$$

Where $A_{1-2}$ is the contact area between cell 1 and 2. In this formulation, if cell 3 is not contacting cell 1, then $A_{1-3}D_3$ would be zero and thus remove the cell 3 Delta contribution. The constant c is the same as in the two-cell model, and thus the only addition would be the Notch activation from the third cell. This way of introducing OR logic into a Hill function is also used in other studies (Kirouac et al. 2013), and we feel that it appropriately encompasses the contribution of additional cells. The general Notch-Delta dynamics can be written as the following:

$$\frac{dN_i}{dt} = \frac{\left(\sum_{i\neq j} A_{i-j}D_j\right)^r}{c^r+\left(\sum_{i\neq j} A_{i-j}D_j\right)^r} - N_i \tag{5}$$

$$\frac{dD_i}{dt} = v\left(\frac{1}{1+(\mathrm{N_i}/\mathrm{K_D})^h} - D_i\right) \tag{6}$$

Both c and v are assumed to be equal to one, leaving four total parameters: $K_D$, which we assume to be the same between different cells, and three surface area variables $A_{1-2}$, $A_{2-3}$, and $A_{3-1}$.

### 3 Exploring the behavior of the Three-Cell Model

#### 3.1 Three back-to-back cells

The first situation that we considered is one where three cells are in contact in an end-to-end orientation. If cell 1 is the middle cell, then there is no contact between cell 2 and 3. This means that the parameter $A_{2-3}$ is set to zero. Using this simplification, we assessed the effect of additional cell contact on Notch and Delta levels at equilibrium. To keep these examples comparable to the two-cell model, the equilibrium levels of Notch and Delta are calculated as a function of $K_D$ and $A_{1-2}$ while keeping $A_{1-3}$ fixed (Figure 1). In these examples, setting the $A_{1-3}$ to zero reproduces the two-cell model, and in this limit, $1/A_{1-2}$ is the same as $K_N$ in the two-cell model plots.

What becomes clear in these simulations is that with increasing cell contact of an additional cell (cell 3), the Delta dynamics become less and less dependent on cell 2 contact. For example, at $A_{1-3}$ of 100, cell 1 is going to differentiate as shown by the high Notch state for all variables of $A_{1-2}$ (Figure 1A). Additionally, since cell 1 is in contact with both cell 2 and 3, there is an asymmetric regime where cell 1 differentiates while cell 2 remains stem as shown by high-Delta low-Notch (Figure 2). Cell 3 dynamics are largely independent of cell 2 dynamics since they are not directly contacting one another (Figure 3). In cell 3, if $A_{1-3}$ is very low, the cell stays as stem, and if $A_{1-3}$ is very high, the cell will generally differentiate except for a low $K_D$ regime. The only dependence between cell 2 and cell 3 occurs at $A_{1-3} \approx 1$, where the combined effects of cell 2 and 3 push cell 1 to differentiate.

When looking at Delta levels across all three cells, the only way to increase equilibrium Delta levels in Notch-positive regimes is by increasing $K_D$, which is what the two-cell model also suggests. These simulations show that for the range of parameters tested, increasing the number of cells in the model is not going to induce a high-Delta high-Notch state without an increase in $K_D$. Mathematically speaking, the addition of cells is incorporated via their contact area in the Notch dynamics equation, and the addition of contacting cells is another way of changing $K_N$ of the two-cell model. And as it was shown previously, changing $K_N$ alone cannot reproduce the high-Delta high-Notch state.

Another way of showing the same effect is to keep $K_D$ constant and look at the effect of $A_{1\text{-}2}$ and $A_{1\text{-}3}$ on Notch and Delta levels (Figure 4). In general, for a low $K_D$ value (Figure 4A), a high-Delta high-Notch state is not achievable, while for higher $K_D$ values (Figure 4B-C), not only a high-Delta high-Notch state becomes achievable, a low-Delta high-Notch state is no longer feasible.

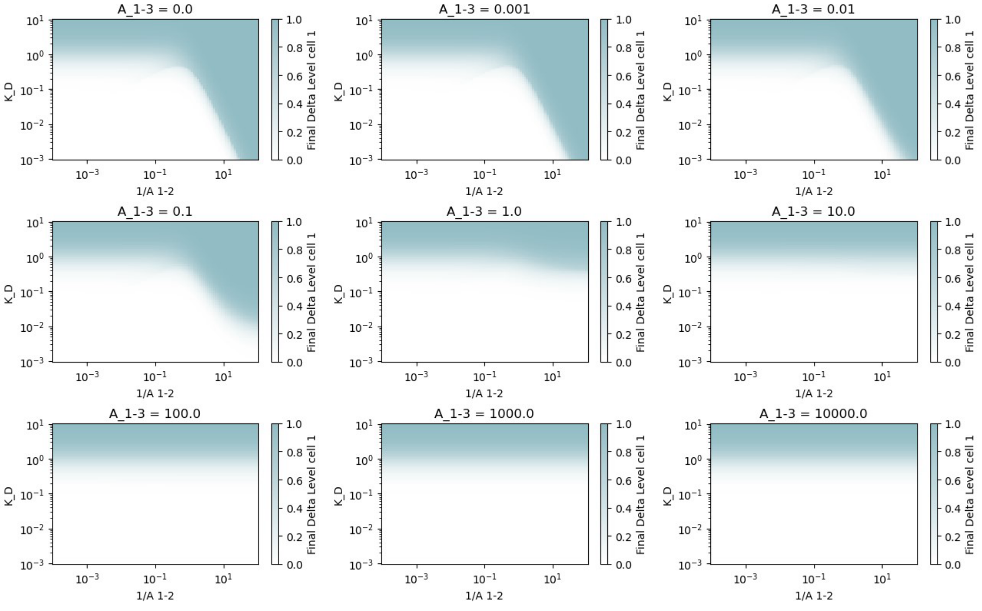


(a)

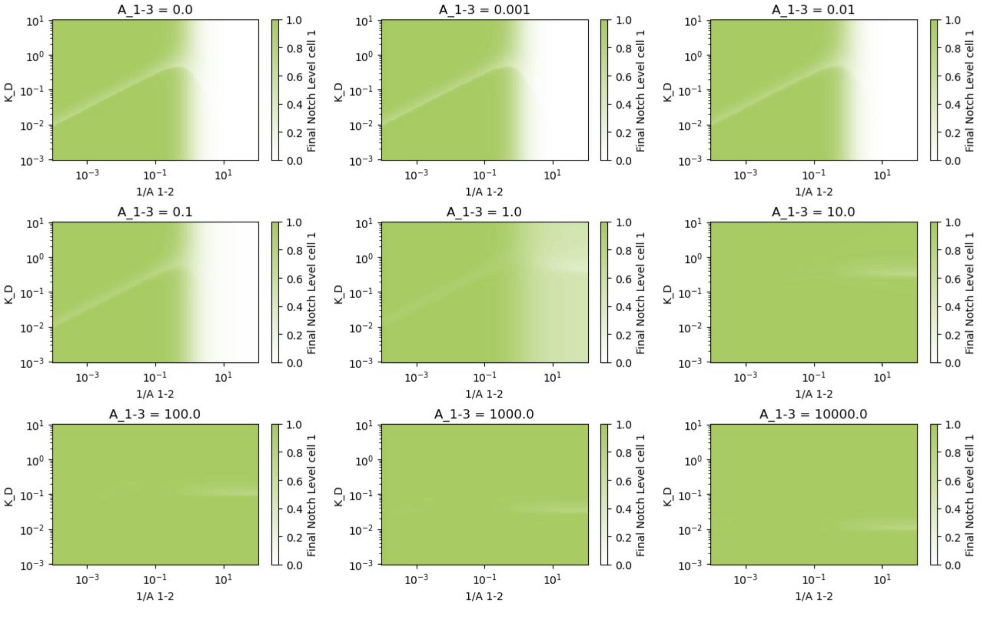


(b)

Figure 1: Equilibrium level of A) Delta and B) Notch as a function of $K_D$ and $A_{1\text{-}2}$ while holding $A_{1\text{-}3}$ fixed for the cell 1 (the middle cell).

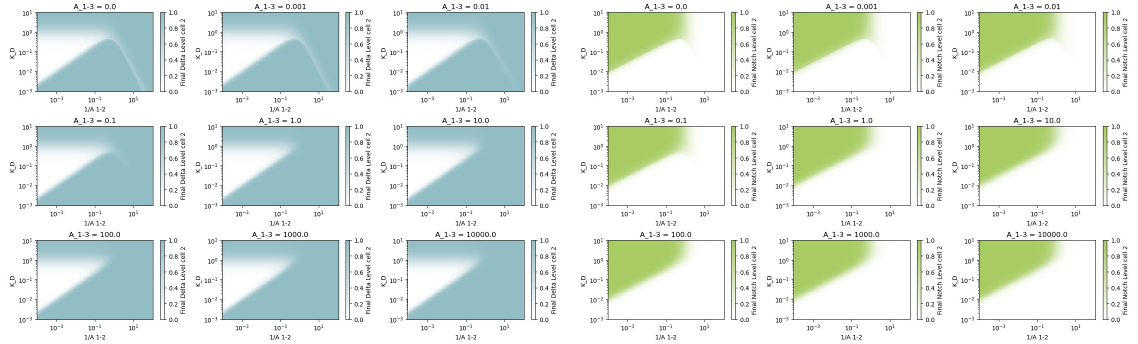


(a) (b)

Figure 2: Equilibrium level of A) Delta and B) Notch as a function of $K_D$ and $A_{1-2}$ while holding $A_{1-3}$ fixed for the cell 2 (one of the side cells).

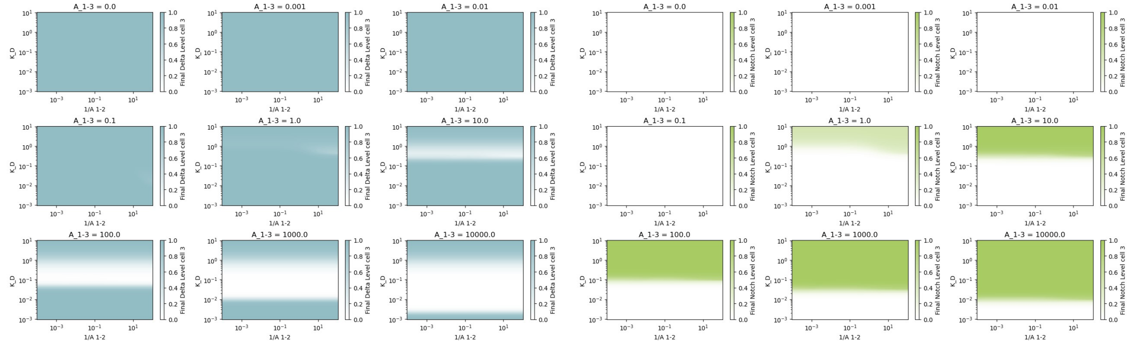


(a) (b)

Figure 3: Equilibrium level of A) Delta and B) Notch as a function of $K_D$ and $A_{1-2}$ while holding $A_{1-3}$ fixed for the cell 3 (the other side cell).

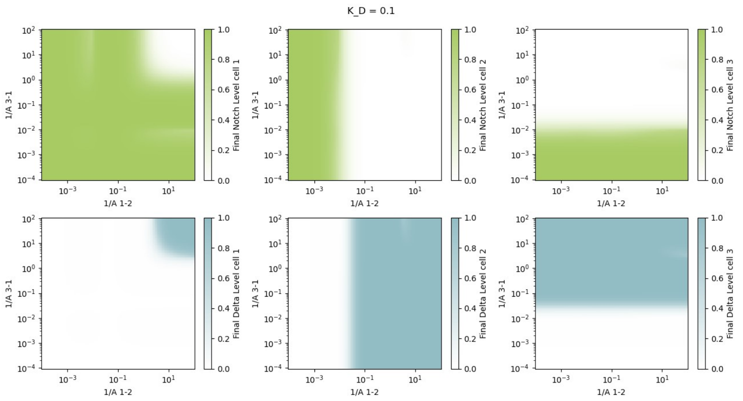


(a)

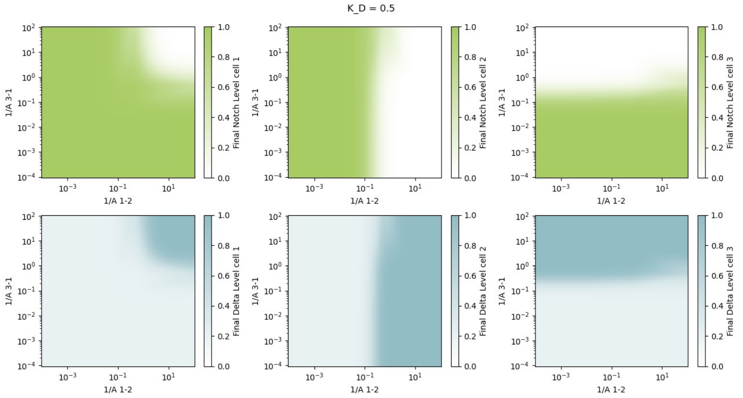


(b)

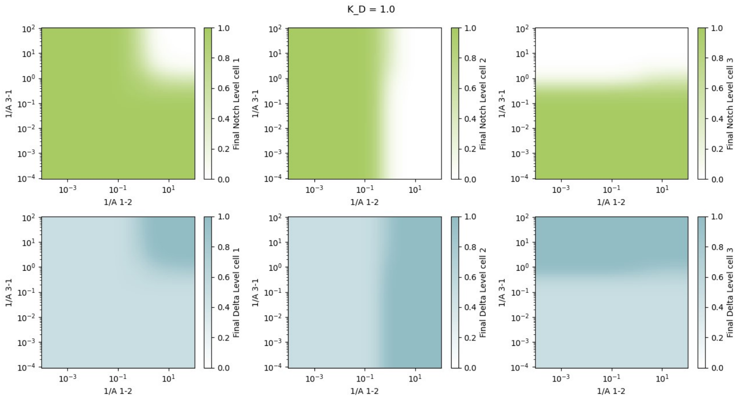


(c)

Figure 4: Equilibrium level of Notch and Delta as a function of $A_{1-2}$ and $A_{1-3}$ for a fixed $K_D$ of a) 0.1, b) 0.5, c) 1.0.

From the two-cell model results, we predicted that the increase in $K_D$ would accompany an increase in Notch activation speed. We wondered whether the cell clustering influenced this. To assess this, we generated the Notch activation speed for cell 1,2, and 3 as a function of $K_D$ and $A_{1-2}$ while keeping $A_{1-3}$ fixed (Figure 5). Quite interestingly, the Delta and Notch equilibrium levels of cell 1 are not affected at low $A_{1-3}$ values of 0.001-0.01 (Figure 1), but Notch activation speed increases in this regime (Figure 5a). This effect can also be shown by fixing $K_D$ and looking at Notch activation speed as a function of $A_{1-2}$ and $A_{1-3}$ (Figure 6). This effect is clearer for $K_D$ of 0.1 and 0.5, before Notch activation speed reaches maximal level due to the increasing $K_D$.

#### 3.2 Extension to the full model

In the previous section, we assumed that the three cells are in a line so that we can remove one contact area parameter from the equation. To ensure that the conclusions from the previous section still apply when all the cells are contacting one another, we considered the full model. The difficulty with this model is the presence of four parameters, which means that to analyze this model, we need to keep at least two parameters fixed. From the examples in the previous section, the $K_D$ values of 0.1, 0.5, and 1.0 are a reasonable range. Thus, we decided to look at cell 1 Notch and Delta levels as well as Notch activation speed as a function of $A_{1-2}$ and $A_{1-3}$ while keeping $A_{2-3}$ fixed at the values of 0.1, 1.0, and 10 (Figure 7). Similar to before, for the range of different contact area parameters tested, there is no high-Delta high-Notch state without increasing $K_D$, emphasizing how the perturbation to $K_D$ is necessary to produce Delta-positive Notch-positive cells. Additionally, as in the previous section, the increase in contact area can result in higher Notch activation speed, generalizing the conclusions from the prior section.

#### 3.3 Conclusions

From studying the model of three cells interacting with each other, we see that increasing cell-cell contact cannot generate a high-Delta high-Notch state similar to the Delta-positive Notch-positive cells observed in intestinal stem cells after injury. The addition of another cell only effectively changes the $K_N$ of the two-cell model, and this change is not sufficient to induce a high-Delta high-Notch state. However, the addition of other cells can increase the speed of Notch activation. Since increasing $K_D$ also causes this effect, the combined effect of increasing $K_D$ and cell clustering would further increase the Notch activation speed and help with tissue regeneration.

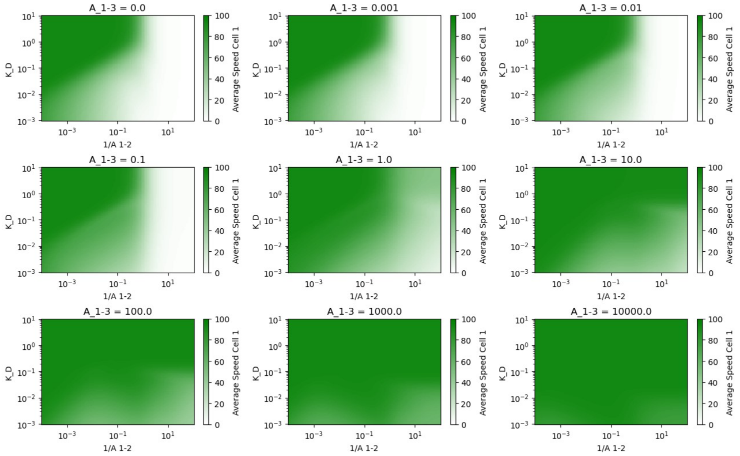


(a)

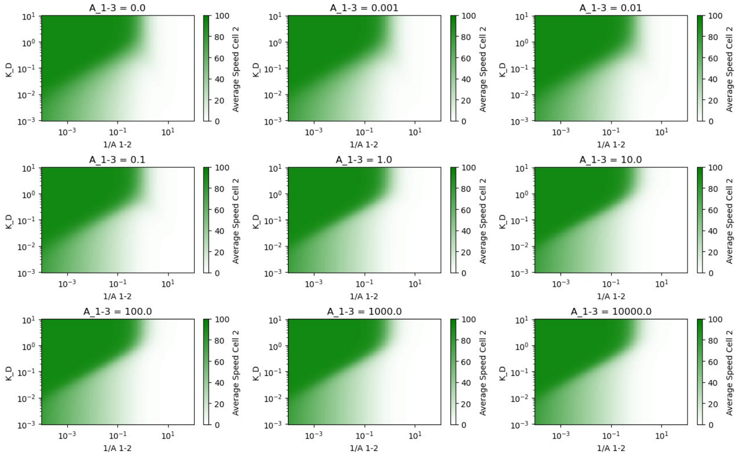


(b)

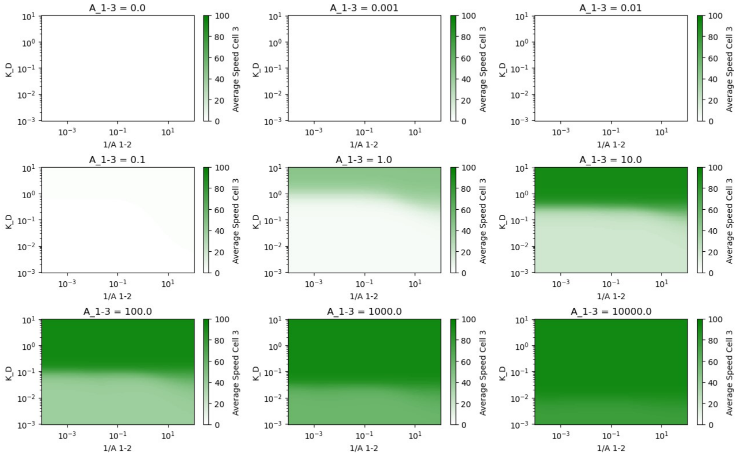


(c)

Figure 5: Notch activation speed as a function of $K_D$ and $A_{1-2}$ while keeping $A_{1-3}$ fixed for a) cell 1, b) cell 2, and c) cell 3.

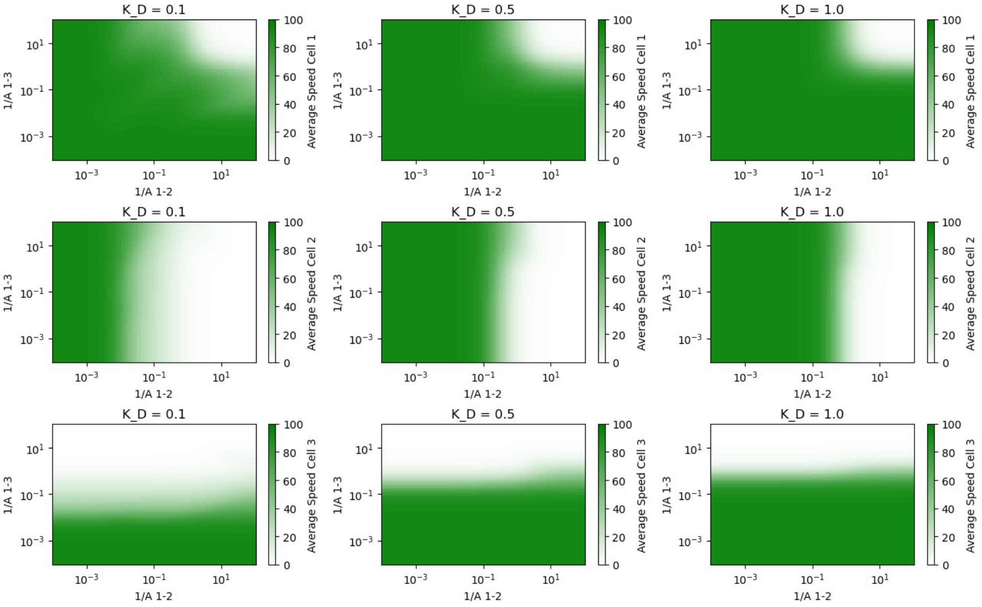


(a)

Figure 6: Notch activation speed as a function of $A_{1-2}$ and $A_{1-2}$ while keeping $K_D$ fixed for cell 1, 2, and 3.

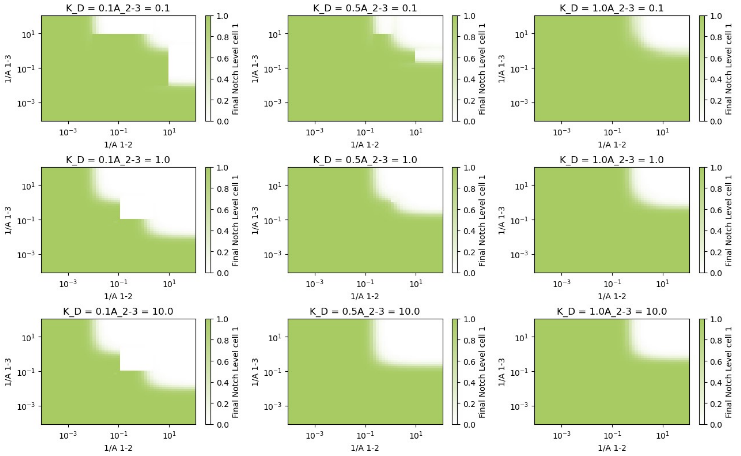


(a)

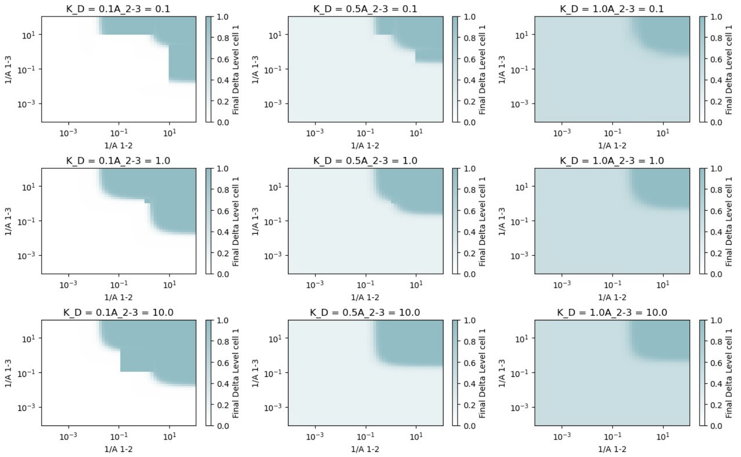


(b)

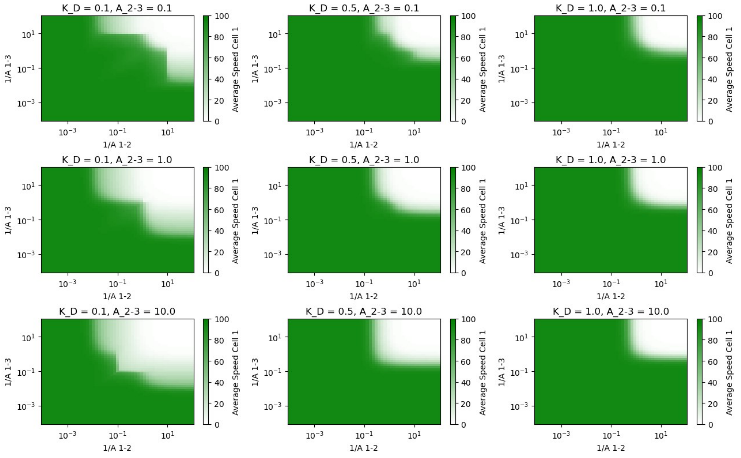


(c)

Figure 7: The equilibrium levels of a) Notch, b) Delta, and c) Notch activation speed as a function of $A_{1-2}$ and $A_{1-3}$ while keeping $K_D$ and $A_{2-3}$ fixed for cell 1.
